# Architectural Fragmentation and Immune Reprogramming Define Early Malignant Transition in Lung Adenocarcinoma

**DOI:** 10.1101/2025.10.23.684213

**Authors:** Beibei Huang, Bo Zhu

## Abstract

The earliest steps of lung adenocarcinoma (LUAD) evolution remain poorly resolved because existing spatial-omics technologies cannot scale to the large archival cohorts required to capture rare transitional states. Here, we reconstructed a stage-resolved spatial atlas of LUAD progression using ROSIE, a deep-learning framework inferring 50-plex protein expression from routine H&E slides (114 specimens; 3.8 million cells). Using a pathology-defined precursor continuum (Normal→AAH→AIS→MIA→IAC) as a pseudotemporal axis, this scalable virtual multiplexing approach reveals a previously unrecognized two-phase immune reprogramming cascade tightly coupled to a measurable structural transition we term architectural fragmentation. During the AAH→AIS transition, transient adaptive immune engagement collapses, marked by a 48.4% loss of CD8 + T cells and the emergence of microfragmented epithelial–stromal interfaces. Subsequent AIS→MIA→IAC transitions are dominated by myeloid-driven immunosuppression, expansion of CD163 + macrophage niches, and the establishment of an immune-excluded architecture. Fragmentation severity in AIS strongly predicts downstream invasive potential, positioning it as an early, quantifiable hallmark of malignant transition. These findings define a mechanistic sequence linking structural disintegration, immune collapse, and invasive outgrowth and demonstrate that H&E-based virtual multiplexing enables cohort-scale dissection of early cancer evolution and identification of actionable windows for immunoprevention. The ROSIE predictions were validated against imaging mass cytometry (IMC) across 25 precursor samples, achieving concordance up to r ≥ 0.90 (93% of marker–cell-type pairs at r ≥ 0.80). A complementary comparison against MIPHEI-ViT, an independent virtual multiplexing framework, corroborated the structural and pan-immune components of this signature across all stages, while highlighting fine-grained immune-lineage resolution as a shared open challenge for H&E-based virtual multiplexing.

## Introduction

Lung adenocarcinoma (LUAD) progresses through a precursor continuum, atypical adenomatous hyperplasia (AAH), adenocarcinoma in situ (AIS), and minimally invasive adenocarcinoma (MIA); however, the cellular and spatial mechanisms governing these earliest transitions remain poorly defined [1–7]. Most of the existing knowledge is derived from advanced tumors, leaving the actionable window of early evolution largely unexplored. Technological barriers include high-dimensional spatial profiling platforms such as multiplex immunofluorescence, imaging mass cytometry, and spatial transcriptomics, which provide deep molecular resolution but cannot scale to the large archival cohorts required to capture rare transitional states [8–19]. Computational models built on these assays offer mechanistic insight but remain tied to low-throughput, specialized data. In contrast, H&E slides are ubiquitous; however, conventional analysis lacks the molecular resolution needed to resolve immune composition, functional states, or spatially organized microenvironmental programs [20–23].

To overcome this scalability gap, we applied ROSIE, a deep-learning framework that converts routine H&E slides into 50-marker spatial maps, bypassing the throughput and cost limits of conventional multiplexed imaging. Using ROSIE-derived single-cell maps from 114 LUAD specimens, we show early malignant transition is accompanied by measurable structural and immune shifts, identifying stage-specific vulnerabilities for precision interception. This work makes four contributions: (i) the first quantitative, cohort-scale definition of architectural fragmentation as a structural hallmark measurable at the AIS stage and predictive of invasive potential; (ii) identification of a conserved two-phase immune reprogramming cascade linking early adaptive immune collapse to later myeloid-driven immunosuppression; (iii) validation of ROSIE-derived spatial signal against orthogonal imaging mass cytometry ground truth; and (iv) an independent cross-model comparison against MIPHEI-ViT corroborating the structural and pan-immune components while noting fine immune-lineage resolution as a shared challenge. An overview of this coordinated remodeling across the LUAD continuum is shown in Fig. 1.

**Fig. 1.**
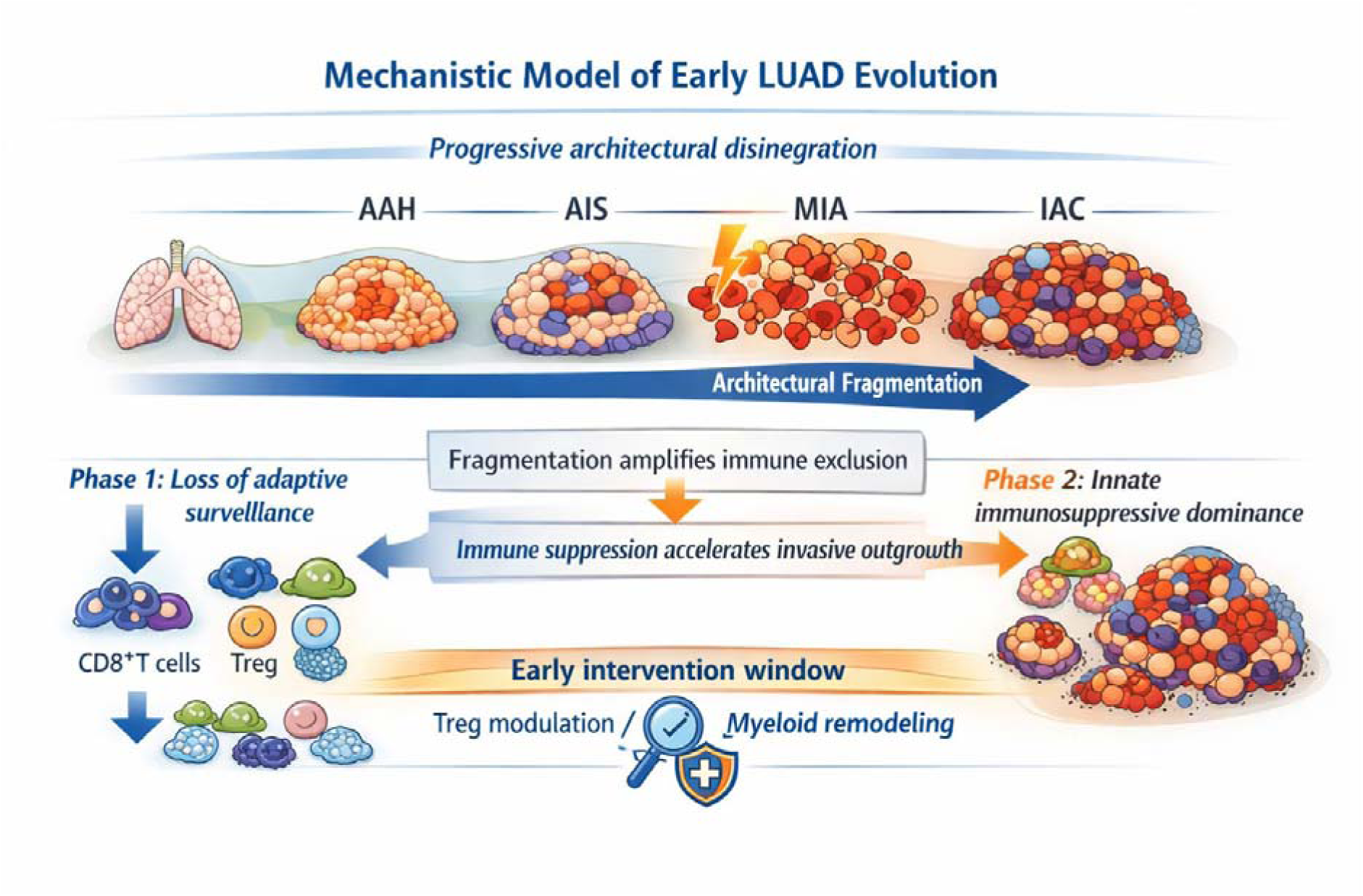
Mechanistic model of early LUAD evolution. Schematic summary illustrating coordinated immune–architectural remodeling during lung adenocarcinoma progression. The upper panel depicts progressive architectural disintegration from cohesive alveolar structures (Normal,JAAH,JAIS) to fragmented and invasive clusters (MIA,JIAC). The lower panel shows a two-phase immune cascade: early adaptive collapse marked by CD8⁺JT-cell depletion and endothelial loss, followed by myeloid-driven immunosuppressive dominance with macrophage expansion. Bidirectional arrows highlight reciprocal reinforcement between structural fragmentation and immune exclusion. The AIS–MIA boundary represents the inflection point of malignant transition and defines an early intervention window for Treg modulation or myeloid remodeling.

## Materials and Methods

### Study design and inference of progression

This study is based on cross-sectional tissue specimens rather than longitudinal follow-up, and therefore all uses of terms such as “progression,” “transition,” and “cascade” refer to a pathology-defined precursor continuum rather than true temporal evolution within individual patients. We treated the sequence Normal→AAH→AIS→MIA→IAC as a well-established biological continuum and used this ordering as a pseudotemporal axis to compare structural and immune features across stages.

To reduce potential biases introduced by interpatient variability inherent to cross-sectional designs, we included 33 patients who harbored multiple precursor lesions (AAH, AIS, and/or MIA) at the same time point. Paired analysis of these multifocal lesions showed that within-patient differences mirrored cohort-level stage trends, supporting the interpretation that observed stage-resolved patterns reflect a conserved biological sequence rather than population-level artifacts.

### Human samples, histopathology, and whole-slide imaging

H&E-stained whole-slide images (WSIs) were obtained from a retrospective cohort of 114 human lung tissue samples spanning the evolutionary continuum of LUAD, including normal lungs, AAH, AIS, MIA, and invasive adenocarcinoma (IAC). All samples were collected under protocols approved by the Institutional Review Board (IRB) and adhered to the ethical guidelines for human subject research. Cases were included if they had a confirmed histopathological diagnosis at one of the five defined stages and sufficient tissue quality for WSI digitization. Cases were excluded if the slides showed significant artifacts, insufficient tissue area (<1 mm²), or ambiguous staging diagnosis upon consensus review.

The cohort comprised 62 patients with 114 lesions spanning five histopathological stages (40 AAH, 22 AIS, 18 MIA, 34 IAC, and 10 normal lung tissues; median age, 71 years; range, 53–82 years; 38 female, 24 male). Thirty-three patients presented with multiple synchronous lesions. Histopathological diagnosis and staging were performed by board-certified pathologists according to IASLC/ATS/ERS classification criteria [24]. The WSIs were digitized at 40× magnification (∼0.25 µm/pixel). A total of 816 representative regions of interest (ROIs) were selected to encompass diverse epithelial, stromal, and immune compartments across disease stages.

### ROSIE model and virtual multiplex immunofluorescence inference

Biomarker inference was performed using the pretrained ROSIE (RObust in Silico Immunofluorescence) framework [25]. ROSIE is a deep learning model that translates H&E-stained morphological features into high-fidelity virtual multiplex immunofluorescence (v-mIF) signal intensities, originally developed and benchmarked on a large-scale, multi-institutional dataset of paired H&E and mIF images. The pretrained model was applied as a frozen inference engine to the independent LUAD precursor cohort, ensuring that downstream spatial analyses reflect biological generalization across disease stages rather than overfitting the training distribution.

### Comparison with an independent virtual multiplexing model (MIPHEI-ViT)

A limitation of relying on a single virtual multiplexing framework is that mechanistic conclusions drawn from its outputs could in principle reflect model-specific biases rather than generalizable biology. To address this, we additionally applied MIPHEI-ViT [26], a publicly released Vision Transformer-based model for H&E-to-multiplex immunofluorescence prediction, as a second, independently developed frozen inference engine. Because MIPHEI-ViT outputs a partially overlapping 16-marker panel rather than the 23-marker IMC panel, this cross-model comparison was performed on an expanded set of 52 ROIs spanning all five stages (Normal = 4, AAH = 6, AIS = 13, MIA = 12, IAC = 17; 203,109 cells total) drawn from the same 114-specimen cohort. MIPHEI-ViT predictions were generated using its released pretrained weights without task-specific fine-tuning. Because ROSIE (∼0.8 µm/pixel) and MIPHEI-ViT (0.5 µm/pixel) were developed and run independently with no shared tiling or coordinate convention, per-ROI spatial registration was performed by template-matching ROSIE’s DAPI channel against MIPHEI-ViT’s Hoechst channel (rescaled 1.6×) via normalized cross-correlation (mean confidence 0.40, range 0.15–0.56); registration accuracy was confirmed by visual inspection of registered cell centroids (Supplementary Fig. S13a). ROSIE’s own per-cell marker calls were treated as the reference signal, since the goal was to test cross-model agreement rather than absolute accuracy against wet-lab ground truth, which was established independently via the IMC comparison above. Concordance for the 16 shared markers was assessed at two spatial resolutions: single-cell level (each cell’s ROSIE signal versus mean MIPHEI-ViT signal within a nucleus-scaled radius of its registered centroid; 203,109 cells), and native-patch level, in which both predictions were block-averaged onto ROSIE’s intrinsic ∼64-pixel inference grid to remove the resolution mismatch between ROSIE’s patch-wise output and MIPHEI-ViT’s dense pixel-wise output (19,202 matched patches).

### Single-cell segmentation and feature extraction

The standardized 50-marker feature vectors (n = 3,817,675) derived from the current watershed-based framework provided a high-fidelity map of the tumor microenvironment. Future work will explore the integration of Variational Autoencoders (VAEs) and Diffusion Models to transition from deterministic intensity extraction to deep representation learning, enabling a more nuanced identification of latent cellular states and dynamics.

### Cell type classification and dimensionality reduction

The segmented cells were categorized into 14 distinct phenotypic classes using a hierarchical classification strategy based on canonical lineage markers. A fuzzy classification algorithm (described in **Supplementary Fig. S11**) evaluated the co-expression of structural and immunological proteins (e.g., pan-cytokeratin for epithelial cells, CD3/CD4/CD8 for T-cell subsets, and CD68/CD163 for myeloid lineages); detailed marker combinations and thresholds are provided in Supplementary Table S2. The UMAP embeddings were generated from the normalized 50-marker feature vectors. All dimensionality reduction and clustering analyses were implemented in Python (v3.9) using Scanpy (RRID:SCR_018139) and UMAP (RRID:SCR_018217).

### Spatial clustering and architectural quantification

Density-Based Spatial Clustering of Applications with Noise (DBSCAN) was applied to the Cartesian coordinates (x, y) of single-cell centroids. The coordinates were z-score normalized prior to clustering, and the neighborhood search radius (ε) was stage-specifically optimized (ranging from 0.003 to 0.01) to account for context-dependent cellular densities. For each histopathological stage, we derived the following quantitative architectural metrics: cluster enumeration, size distribution, macrocluster fraction, and spatial noise proportion. The robustness of the structural findings was confirmed through parameter sensitivity analysis.

### Population dynamics modeling using timed Petri nets

Timed Petri nets are well suited for discrete, stage-defined systems, where continuous pseudotime cannot be inferred. This formalism was chosen over single-cell trajectory inference methods such as Monocle or PAGA because those methods assume a continuous transcriptomic manifold reconstructed from individual cells, whereas the present data comprise discrete, pathology-defined population states measured at the protein level across cross-sectional specimens; timed Petri nets instead quantify directional population flux between defined stages without requiring pseudotime assumptions or single-cell resolution. Stage-ordered cellular trajectories were reconstructed using a Timed Petri Net (TPN) framework. Places represent the five discrete histopathological stages, with markings corresponding to the observed cell type abundances. Transitions represent biological shifts between successive stages, with magnitudes derived from empirical log-fold changes in cell abundance. Structural robustness was assessed by introducing ±10% stochastic variance into the input cell abundance across 1,000 simulation runs.

### Pathway activity inference and niche-level analysis

Pathway activity scores were computed using weighted marker combinations derived from the ROSIE-predicted protein intensities. Functional signatures, including cell cycle progression, T cell exhaustion, and stromal signaling, were defined by aggregating the normalized intensities of biologically relevant markers (Supplementary Table S2). The analysis was performed at the global tissue level (per ROI) and discrete cellular niches (DBSCAN-defined neighborhoods). Variability across stages was quantified using Kruskal-Wallis tests followed by post-hoc Dunn’s test with Bonferroni correction (n = 114 specimens across five stages). Statistical analyses were performed using SciPy (v1.9, RRID:SCR_008058) and Scikit-learn (RRID:SCR_002577).

## Results

### ROSIE enables scalable multiplex biomarker inference from routine H&E slides

To resolve early microenvironmental transitions at single-cell resolution, we applied ROSIE to generate v-mIF predictions directly from H&Estained tissue sections. By leveraging a cohort of whole-slide images, ROSIE predicted the expression levels for a comprehensive 50-biomarker panel across 816 ROIs, encompassing 3,817,675 individual cells spanning the full LUAD evolutionary continuum.

A key advantage of ROSIE is its ability to perform progressive annotation enrichment using expanded biomarker panels. While a baseline 10-biomarker panel (e.g., PD-L1, CD8, PanCK, FoxP3) captures primary immune and epithelial compartments, the expansion of 50 biomarkers integrates immune checkpoint molecules (PD-1, LAG-3, TIGIT, VISTA, and ICOS), myeloid-lineage markers (CD68, CD14, HLA-DR, Gal3, MPO), B-cell regulators (CD20, CD21, CD38, CD79a), and functional metabolic/antigen-presentation markers (HLA-A, HLA-E, IFN-γ, IDO1). This scalability allows the sequential generation of v-mIF maps with increasing complexity, without requiring additional tissue or staining (Fig. 2).

**Fig. 2.**
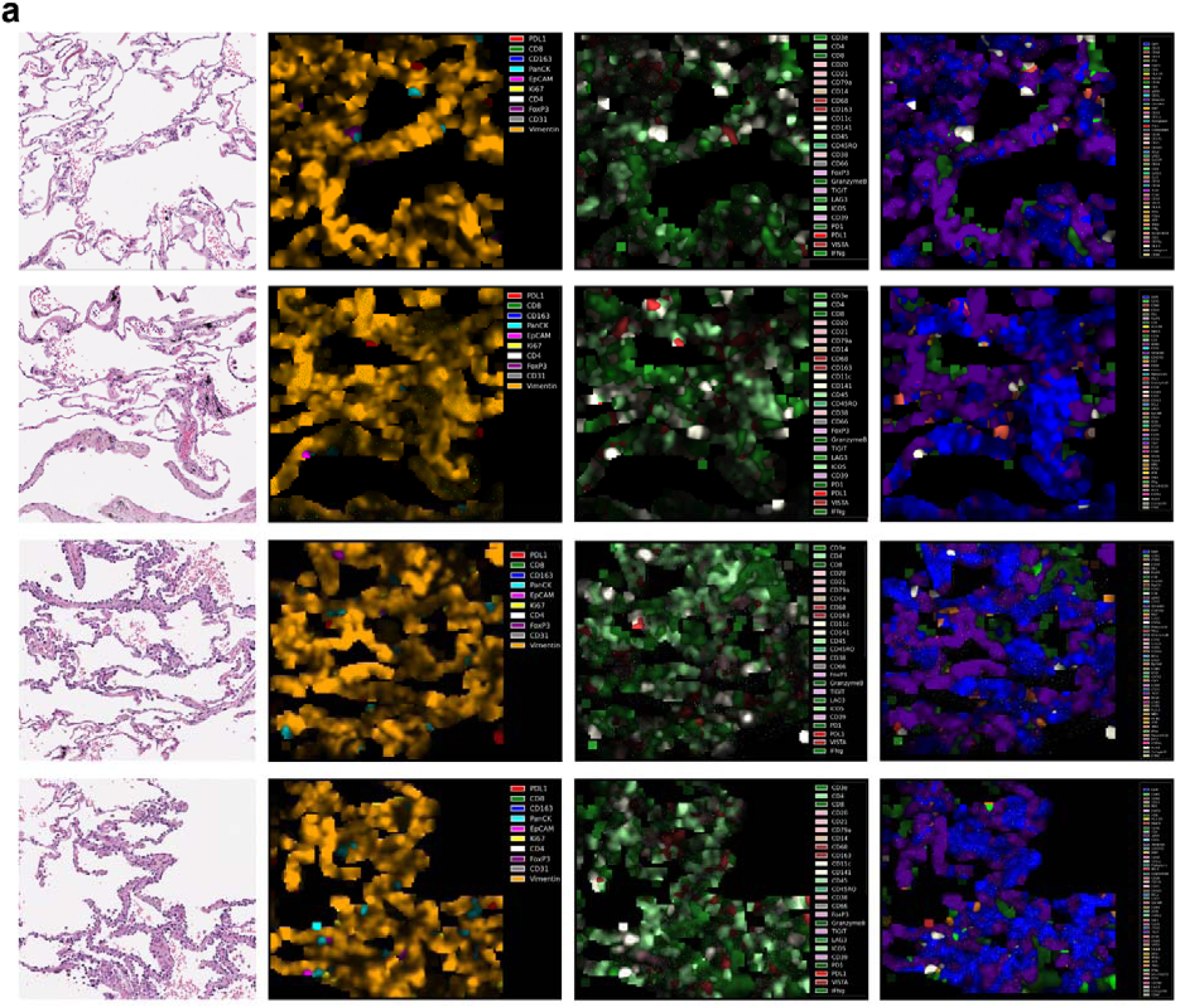
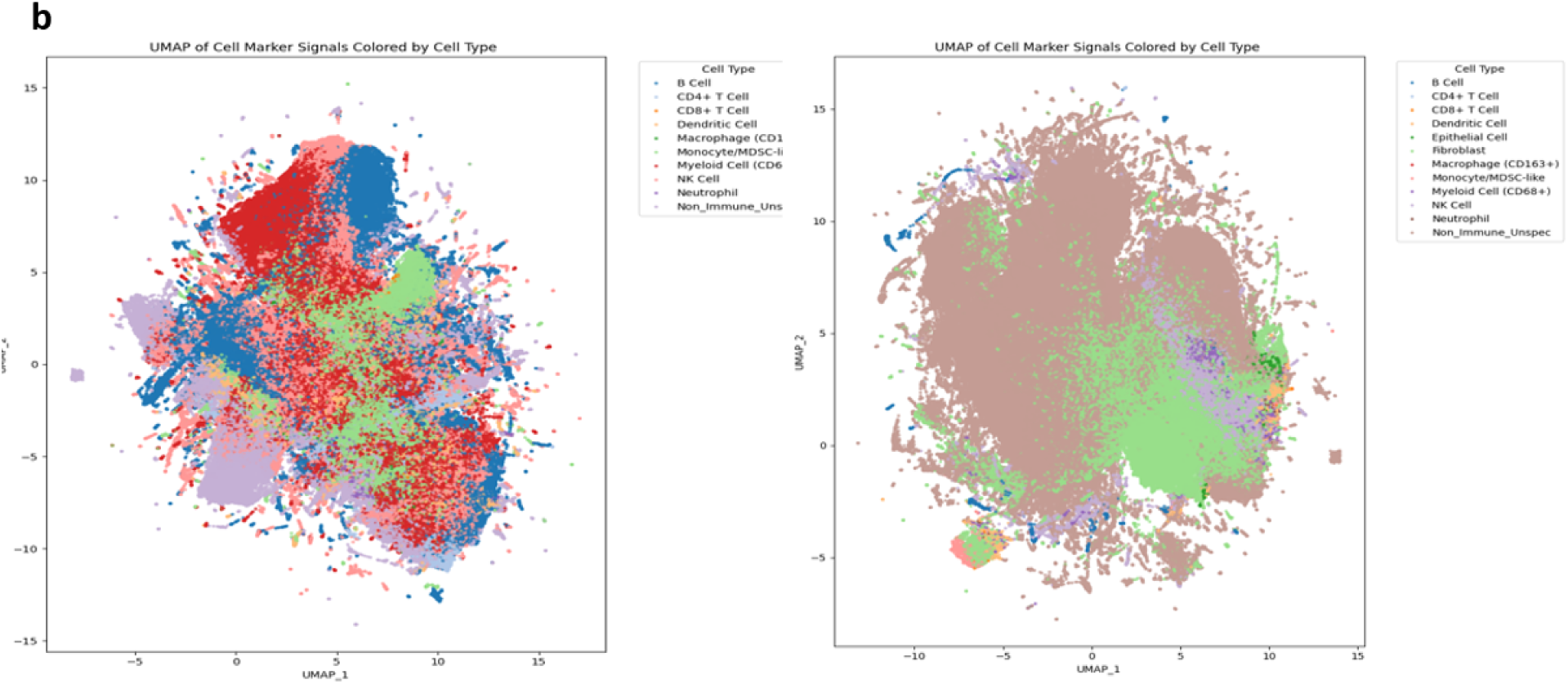
Scalable virtual multiplexing via ROSIE enhances spatial annotation and immune lineage resolution. (a) Each row represents an independent tissue ROI showing ROSIE-generated v-mIF predictions using 10-, 25-, and 50-biomarker panels from the same H&E input. As panel size increases, previously unannotated regions become classified (increased color diversity). White arrows highlight newly annotated regions resolved by the expanded panel. Scale bar: 100 µm. (b) UMAP visualization comparing 25-marker versus 50-marker panels across 3,817,675 cells colored by cell type. The 50-marker panel resolves eosinophils and mast cells absent in the reduced panel due to missing lineage-specific markers (GATA3, MPO, Galectin-3).

UMAP-based dimensionality reduction on predicted marker signals demonstrated that the 50-marker panel revealed significantly finer cellular heterogeneity than the 25-marker panel, enabling high-confidence identification of populations such as eosinophils and mast cell lineages that remained unresolved in the smaller panel due to the absence of specific markers, including GATA3 and Galectin-3 (Fig. 2b).

### Comprehensive cell phenotyping reveals 14 immune and stromal populations across LUAD progression

Using the 50-marker panel, cells were resolved into 14 distinct functional populations based on canonical lineage markers, including T-cell subsets (CD4 +, CD8 +, Tregs), B cells, NK cells, diverse myeloid compartments (macrophages, dendritic cells, monocytes, and MDSCs), and structural components (epithelial, endothelial cells, and fibroblasts) (Fig. 3, left). Inferred marker expression profiles demonstrated high fidelity to the expected biological patterns: co-expression of CD3 and CD8 in cytotoxic T cells, CD163 enrichment in M2-polarized macrophages, and pan-cytokeratin dominance in epithelial compartments.

**Fig. 3.**
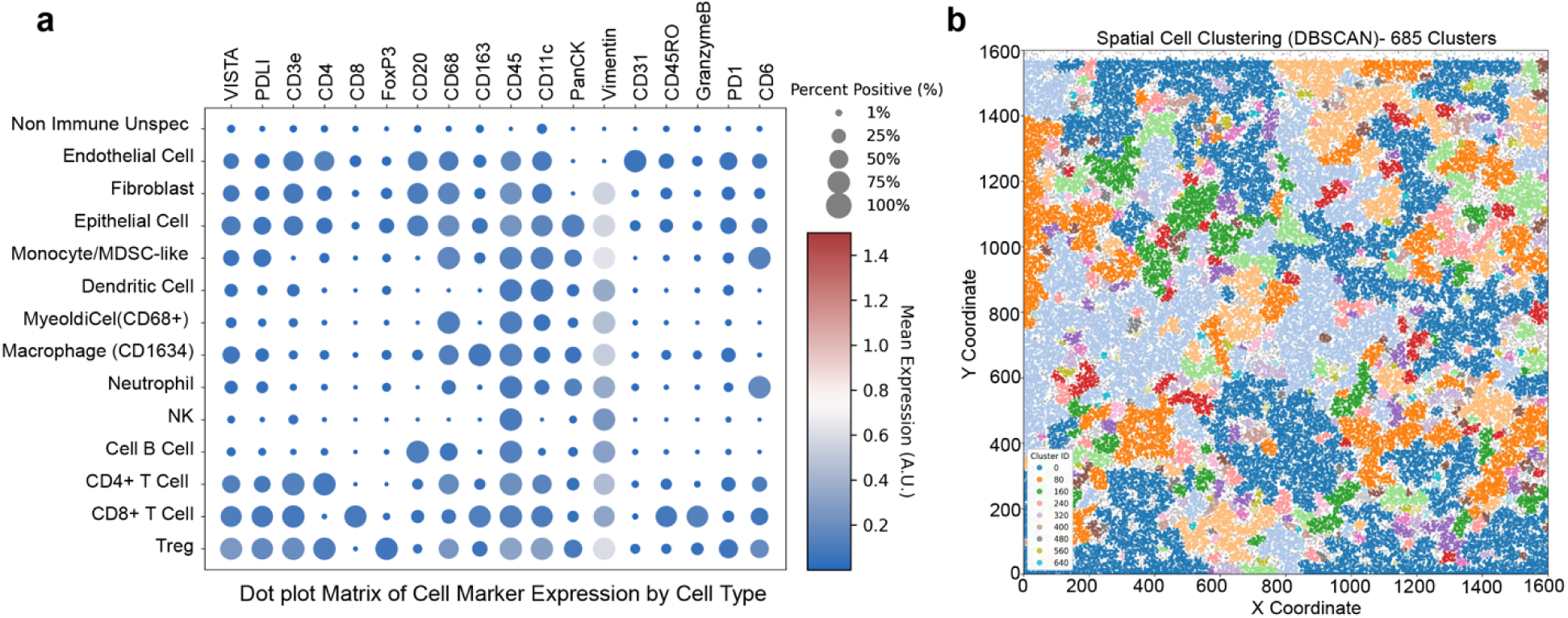
Cell phenotyping and spatial clustering of the LUAD tissue microenvironment. Left: Dot plot showing marker expression across 14 cell types from 3,817,675 cells. Dot size represents percentage of marker-positive cells (1–100%); color intensity indicates mean expression (0–1.4 AU). Right: Spatial distribution of 300,000 sampled cells colored by DBSCAN clusters (685 clusters for downsampled visualization). Gray dots indicate 18,893 unclustered cells.

DBSCAN spatial clustering identified 6,522 microenvironmental niches across the cohort, confirming that ROSIE preserved both the molecular identity and underlying tissue architecture (Fig. 3, right). Detailed cell–cell interaction networks derived from these niche across the evolutionary continuum are provided in Supplementary Figures S1–S5.

### Spatial architecture transitions from cohesive to fragmented during malignant evolution

DBSCAN-based clustering revealed progressive architectural remodeling across the disease stages (Fig. 4). Normal lung tissue (233,459 cells; ε = 0.01) displayed 991 loosely distributed spatial clusters, consistent with the preserved tissue architecture. AAH (511,157 cells; ε = 0.0065) yielded 9,062 clusters with an increased cellular density and defined spatial domains. The AIS samples (437,918 cells; ε = 0.006) exhibited architectural condensation with 12,841 clusters. MIA (776,472 cells; ε = 0.0055) marked the onset of structural fragmentation with 4,936 clusters, whereas IAC (1,858,669 cells; ε = 0.003) demonstrated extensive spatial heterogeneity across 7,314 dense microclusters.

**Fig. 4.**
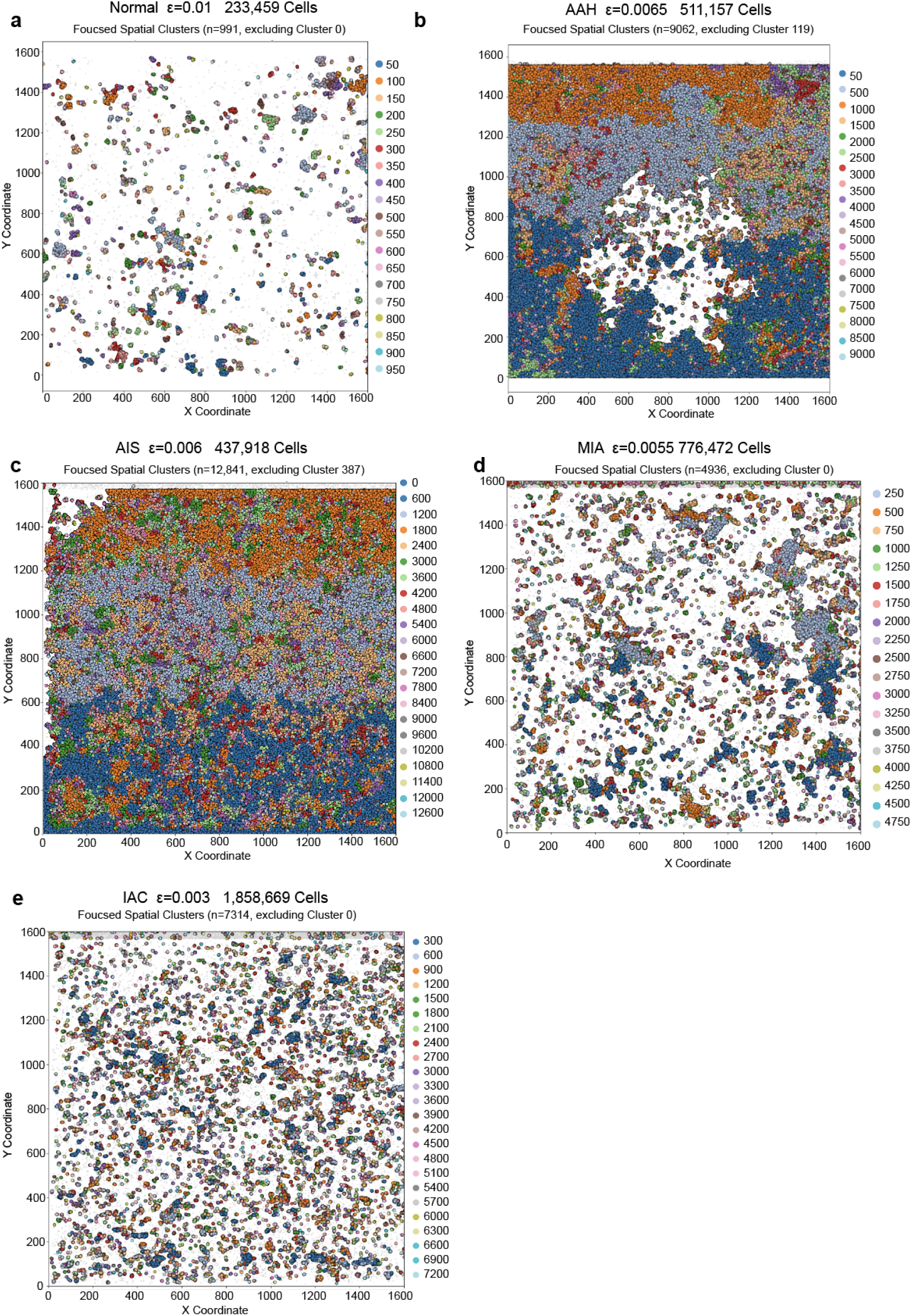
DBSCAN spatial clustering reveals progressive architectural disruption during LUAD development. Spatial clustering maps across Normal lung, AAH, AIS, MIA, and IAC demonstrate distinct patterns of tissue organization during malignant progression. Normal lung displays loosely distributed spatial domains. AAH exhibits increased cellular density with defined spatial domains. AIS shows architectural condensation. MIA marks transitional architecture with emergence of micro-fragmentation. IAC demonstrates dramatic architectural fragmentation with total loss of organized tissue structure. Each point represents an individual cell colored by its DBSCAN-assigned spatial cluster.

This progressive spatial disintegration captures the architectural collapse accompanying malignant progression, reflecting a transition from an organized, immune-surveilled state to a chaotic, invasive landscape. Architectural fragmentation thereby emerges as a quantifiable structural hallmark of malignant transition and potential early biomarker for risk stratification (Table 1). Notably, AIS is the structural inflection point of this trajectory: it shows maximal architectural cohesion (a single dominant macrostructure of 12,841 clusters) immediately preceding fragmentation onset in MIA, positioning cohesion within individual AIS lesions as a candidate quantitative correlate of downstream invasive transition.

**Table 1.**
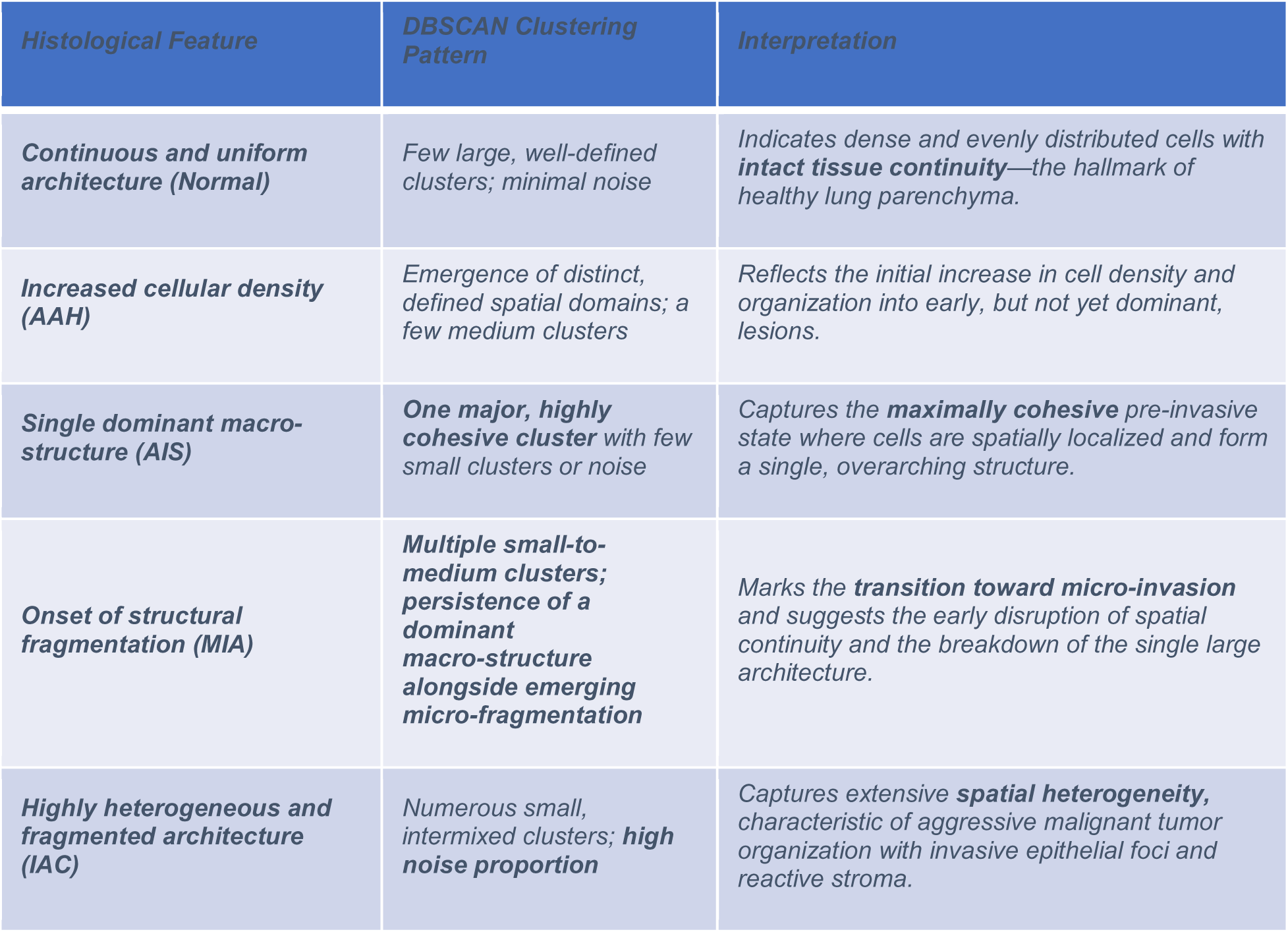
Relationship between histological features and DBSCAN clustering patterns. This table summarizes how distinct histological architectures across lung tissue stages correlate with DBSCAN-derived spatial clustering patterns, linking structural phenotype to clustering behavior and biological significance.

### Immune landscape remodeling reveals coordinated, stage-specific shifts across the precursor continuum

A timed Petri net framework was applied to model population dynamics across the five histopathological stages, revealing coordinated stage-specific remodeling of the immune, stromal, and vascular compartments (Fig. 5a). Trajectory analysis resolved distinct transition patterns across the cellular compartments (Fig. 5b).

**Fig. 5.**
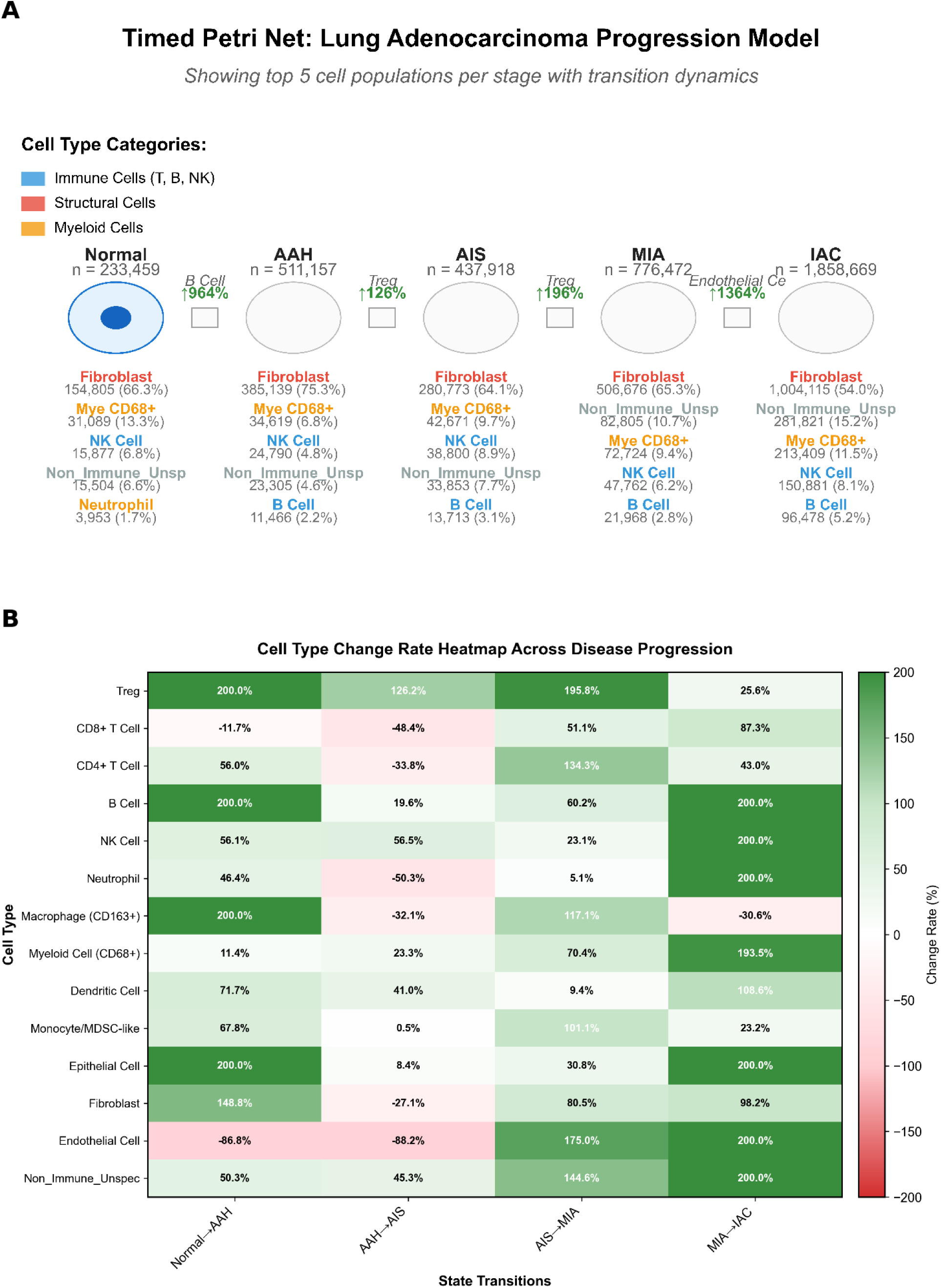
Timed Petri Net model and heatmap of cell population dynamics across LUAD progression stages. (a) The model depicts temporal evolution of cellular compositions through five histopathological states: Normal→AAH→AIS→MIA→IAC. Each circle represents a disease state. Cell types are color-coded by functional category: blue (immune cells), red (structural cells), and orange (myeloid cells). Transition boxes (t1–t4) annotate the most significantly changing cell type and its percentage change. (b) Heatmap showing percentage change in 14 cell populations across four disease transitions. Color intensity indicates magnitude and direction (green: increase; red: decrease). Values capped at ±200%.

Adaptive immunity: CD8 + T cells underwent marked depletion during the AAH→AIS transition (−48.4%) before recovering in later stages. Regulatory T cells (Tregs) expand steadily through mid-stage transitions (up to +200%). B cells infiltrated in two distinct waves (normal →AAH and MIA→IAC), while CD4 + T cells displayed variable dynamics, including initial expansion, transient depletion at AAH→AIS, and subsequent rebound. Myeloid compartment: CD163□ macrophages peaked early before contracting, whereas CD68 + myeloid cells expanded dramatically during MIA→IAC (+193.5%). NK cells accumulated consistently, peaking during MIA→IAC.

Structural populations: Fibroblasts were progressively enriched across all transitions, reflecting sustained stromal remodeling. Epithelial cells showed biphasic expansion consistent with early hyperplasia and late invasive growth. Endothelial cells declined sharply in the early stages, but underwent angiogenic activation beginning at AIS→MIA (+175%), marking the onset of vascularized disease.

Together, these patterns reveal critical inflection points in TME evolution, defining a conserved two-phase immune trajectory: early adaptive engagement, followed by myeloid-driven immunosuppressive dominance.

### Validation of ROSIE-predicted protein expression using IMC ground truth

To assess the fidelity of ROSIE’s virtual multiplexing predictions, we performed targeted validation using a subset of LUAD precursor samples with available imaging mass cytometry (IMC) intensity data [4]. IMC-derived marker intensities from 25 samples spanning five LUAD progression stages were matched to ROSIE-predicted expression values across spatially defined ROIs (n = 15; three ROIs per stage).

Cell-type-stratified correlation analysis of 23 overlapping markers encompassed 74,896 cells across four compartments (Non-Immune unspecified, epithelial, B Cell, and Myeloid/CD68□). Because the IMC reference dataset and ROSIE operate on partially non-overlapping marker sets, correlations were computed between IMC-measured intensities and the ROSIE-predicted signal most biologically proximal within the same cell-type compartment, assessing stage-level trend concordance rather than direct quantitative equivalence. Within this framework, 28 of 30 marker-cell-type pairs (93%) showed concordant directional trends across stages (Pearson’s r ≥ 0.80, p < 0.05), and 23 of 30 (77%) reached r ≥ 0.90, with the strongest agreement in epithelial and myeloid compartments.

To test whether these concordance patterns are specific to ROSIE or generalize across independently developed frameworks, we performed a complementary cross-model comparison against MIPHEI-ViT across 52 ROIs spanning all five stages (203,109 cells; see Materials and Methods). At single-cell resolution, concordance for most of the 16 shared markers was modest (pooled Pearson r range, −0.15 to 0.27), consistent with the resolution mismatch between ROSIE’s patch-wise inference and MIPHEI-ViT’s dense per-pixel output rather than genuine model disagreement or registration error (mean registration confidence 0.40; Supplementary Fig. S13). At ROSIE’s native patch resolution (19,202 matched patches), concordance improved for nearly every marker, most markedly for DAPI (pooled r, 0.18→0.66) and CD45 (pooled r, 0.27→0.35), and remained strong and stage-invariant for DAPI throughout (r = 0.69–0.79 in every stage; Table 2), independently corroborating the structural and general-immune components of the phenotypic signature described above.

**Table 2.** Cross-model concordance between ROSIE and MIPHEI-ViT for representative markers, by disease stage. Values are patch-level Pearson correlation coefficients (52 ROIs; 19,202 matched patches) between ROSIE- and MIPHEI-ViT-predicted signal intensities for the same marker, computed independently within each histopathological stage. DAPI and CD45 (structural/pan-immune) show consistently positive, stage-invariant concordance; PanCK, E-cadherin, and aSMA (epithelial/stromal) show concordance that weakens toward IAC; CD8 (fine immune-lineage) remains weak or absent in every stage. Correlations for all 16 shared markers are provided in the associated data repository (see Data Availability).

| *Marker* | *Class* | *Normal* | *AAH* | *AIS* | *MIA* | *IAC* |
| --- | --- | --- | --- | --- | --- | --- |
| **DAPI** | *Structural (nuclear)* | *0.74* | *0.74* | *0.79* | *0.77* | *0.69* |
| **CD45** | *Pan-immune* | *0.41* | *0.28* | *0.21* | *0.47* | *0.18* |
| **PanCK** | *Epithelial* | 0.13 | 0.34 | 0.12 | 0.22 | 0.01 |
| **E-cadherin** | *Epithelial* | 0.13 | 0.26 | 0.11 | 0.33 | -0.05 |
| **aSMA** | *Stromal* | 0.32 | 0.24 | 0.23 | 0.25 | 0.05 |
| **CD8** | *Immune-lineage* | 0.03 | -0.04 | -0.04 | -0.11 | -0.03 |

By contrast, markers defining the specific immune subpopulations central to the two-phase cascade (CD8, CD3e, FoxP3, CD31, CD163) showed weak or near-zero patch-level concordance (|r| ≤ 0.10) in every stage (Table 2), indicating that fine-grained immune-lineage calls are not yet independently reproducible across current H&E-based virtual multiplexing frameworks and should be regarded as ROSIE-specific pending orthogonal validation. Several epithelial/stromal markers (PanCK, E-cadherin, aSMA, CD4, CD68) also showed progressively weaker concordance toward IAC (early-stage-average-minus-IAC Δr of 0.22–0.25), mirroring the fragmentation trajectory and suggesting more heterogeneous, harder-to-model tissue in invasive disease. Taken together, structural and pan-immune signal is reproducible across independently developed architectures, whereas fine-grained immune-lineage calls remain hypothesis-generating pending orthogonal validation.

### Dynamic pathway reprogramming highlights progressive immune quenching and proliferative dominance

Pathway enrichment analysis across the five histological stages revealed coordinated, stage-dependent reprogramming of signaling networks underlying tumor evolution (Fig. 6; full-resolution stage-specific heatmaps in Supplementary Figures S6–S10, S12).

**Fig. 6.**
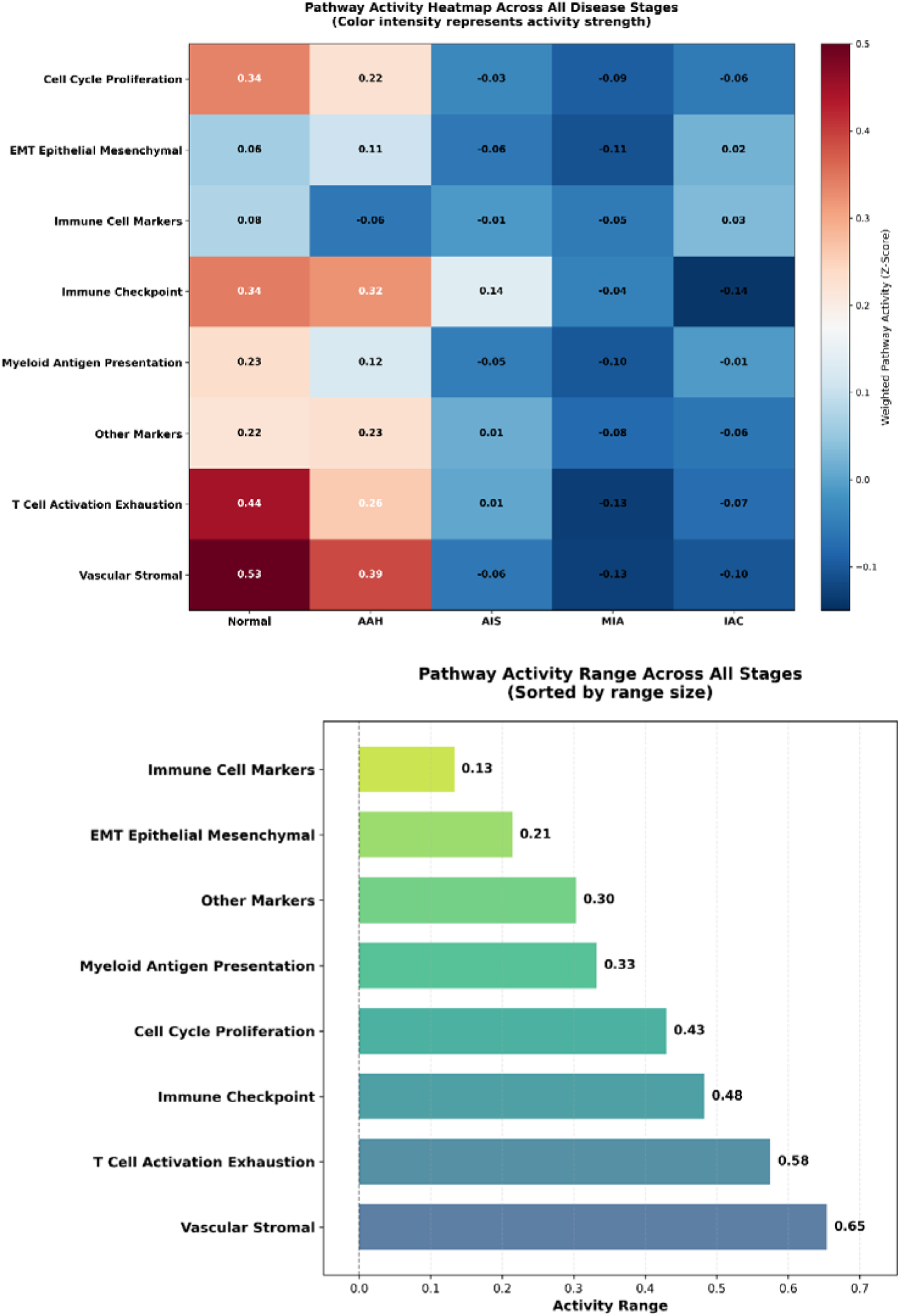
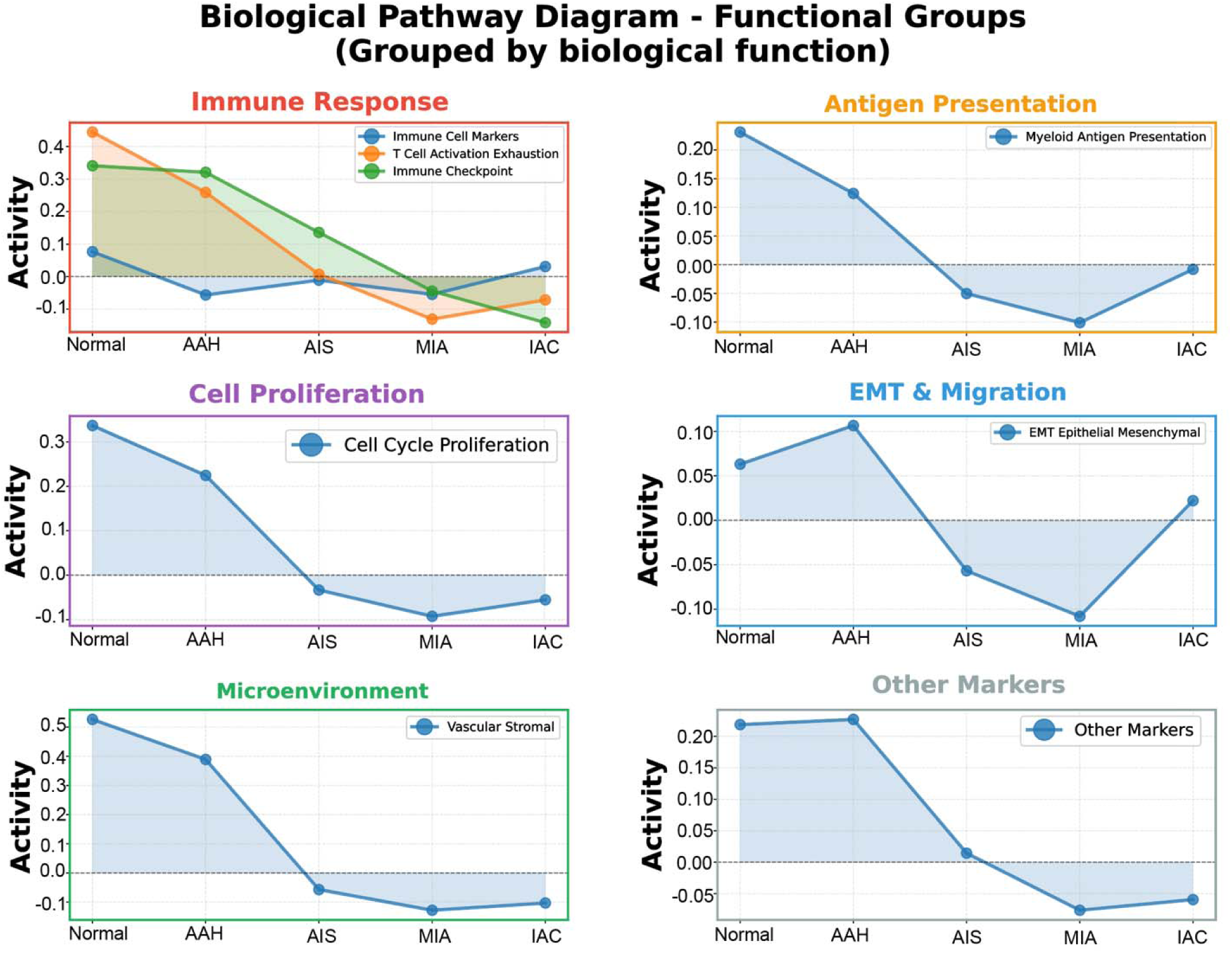
Global pathway activity dynamics across LUAD progression. (a) Normal and AAH show broadly elevated pathway activity in vascular–stromal and adaptive immune programs; advanced stages exhibit coordinated suppression. (b) Pathway variability highlights vascular–stromal and T cell exhaustion pathways as those undergoing the strongest remodeling. (c) Trajectory plots show progressive decline of immune and stromal activities and a biphasic EMT/migratio pattern.

At the global tissue level, Normal and AAH stages showed broadly elevated pathway activity, especially in vascular–stromal and adaptive immune programs, indicating that immune quenching begins during pre-invasive transitions rather than as late events. Advanced stages (AIS–MIA–IAC) exhibited coordinated suppression of immune checkpoints, antigen presentation, and the stromal and vascular pathways (Fig. 6). Niche-level analysis revealed distinct patterns of immune, stromal, and proliferative engagement across individual cell types, with progressive suppression of immune programs concomitant with increased proliferative activity in the epithelial compartments (Supplementary Fig. S12).

Kruskal-Wallis test revealed significant pathway activity variability across histological stages (H(4) = 14.83, p = 0.005), with post-hoc Dunn’s test (Bonferroni-corrected) identifying a significant difference between AAH and IAC (p = 0.006), consistent with the progressive immune quenching trajectory.

## Discussion

ROSIE converts routine H&E slides into high-resolution spatial immunophenotyping maps, thereby overcoming the cost and scalability barriers of conventional multiplex immunofluorescence. Profiling 3,817,675 cells across 816 ROIs from 114 LUAD specimens, we demonstrated that increasing biomarker panel depth progressively improves immune lineage resolution, enabling the detection of rare populations such as eosinophils and mast cells that remain unresolved with smaller panels (Fig. 2). This scalability highlights the potential of computational pathology to extract latent molecular information from archival histology at a scale impractical to antibody-based assays.

Spatial clustering revealed a progressive architectural collapse accompanying malignant progression. Normal lung displayed 991 loosely distributed clusters, reflecting preserved tissue integrity, whereas AAH consolidated into a dominant macrostructure. AIS maintained relative spatial continuity, whereas MIA marked the onset of microfragmentation (4,936 sub-clusters), and IAC showed extensive heterogeneity across 7,314 dense micro-niches (Fig. 4; Table 1). This progressive disintegration positions architectural fragmentation, particularly at the AIS–MIA boundary, as a system-level structural biomarker complementing radiologic and histologic assessments for distinguishing indolent from high-risk lesions.

Beyond its role as a structural biomarker, this fragmentation trajectory invites interpretation through tissue-scale jamming physics, in which dense cellular collectives transition between a solid-like, kinetically arrested state and a fluid-like, freely rearranging state depending on intercellular adhesion, cortical tension, and packing density [27]. This transition has been observed directly in diseased human epithelium [28] and in the collective invasion of carcinoma cells [29,30]. Viewed through this lens, the DBSCAN trajectory in Fig. 4 and Table 1 is descriptively consistent with a jamming-to-unjamming transition: Normal/AAH tissue resembles a kinetically arrested, solid-like state; AIS condenses into a single dominant, highly cohesive macrostructure approaching a percolating cluster; and the MIA→IAC transition, in which this assembly fragments into thousands of microclusters, resembles the breakup of a percolating cluster once local connectivity falls below a critical threshold [31].

Rising pan-immune (CD45□) infiltration across this interval suggests immune influx may lower effective adhesion mechanically as well as immunologically. We offer this framework as an interpretive lens, not a formally demonstrated rigidity transition — the cell shape index p□ was not computed here, though it is directly computable from our segmentation output as a tractable extension.

Spatially resolved cellular analyses revealed a conserved two-phase immune trajectory across the precursor continuum (Fig. 5). Early adaptive engagement characterized the normal-to-AAH transition, with Treg expansion (+200%) and biphasic B cell infiltration, potentially reflecting tertiary lymphoid structure formation. A critical inflection occurred at AAH→AIS, marked by CD8□ T-cell depletion (−48.4%) and early endothelial collapse (−86.8% to −88.2%), indicating immune quenching preceding overt invasion. The AIS→MIA transition was defined by angiogenic reactivation (+175%) and sustained Treg enrichment (+195.8%), whereas MIA→IAC was dominated by myeloid expansion (CD68□ cells +193.5%) and a second B-cell surge (+200%), consistent with immunosuppressive remodeling via IL-10/TGF-β.

Because these mechanistic conclusions were derived from a single virtual multiplexing framework, we sought independent corroboration using MIPHEI-ViT. Cross-model concordance was strong and stage-invariant for nuclear density (DAPI) and general immune infiltration (CD45), supporting the two central claims of this study: that architectural fragmentation is a genuine structural feature of malignant transition, and that immune infiltration undergoes real, quantifiable remodeling across the continuum, rather than artifacts of ROSIE’s training distribution. However, concordance for the fine immune subpopulations at the core of the two-phase cascade (CD8□ T cells, FoxP3 Tregs, CD163□ macrophages, CD3e□ pan-T cells) remained weak in every stage. This is a meaningful boundary on the present findings: the specific quantitative shifts we report for these populations should be interpreted as ROSIE-derived hypotheses that are directionally plausible and internally consistent with the corroborated structural and pan-immune trajectory, but that await confirmation with lineage-specific antibody-based assays before they can be considered independently validated.

To extend this corroboration to the transcriptomic level for the PD-1/PD-L1 checkpoint axis, we cross-referenced the IMC protein trajectory against mRNA-level evidence using a three-tier framework of orthogonal, non-patient-matched sources: an independent human scRNA-seq cohort (GSE164789; Tier 2) and paired mouse IMC plus scRNA-seq in the 129S4-KrasG12D/129S4-Urethane models (Tier 3; Supplementary Fig. S14, S15). CD274 (PD-L1) mRNA declined monotonically across AIS to MIA to IAC in Tier 2 and rose then fell across the mouse continuum in Tier 3, mirroring the corresponding IMC protein trend in both (Tier 3 Spearman rho = 0.8); PDCD1 (PD-1) mRNA reproduced the IMC dip-and-rebound pattern in Tier 2, and rose monotonically with progression in mouse T cells in Tier 3, with no mouse PD-1 protein comparator available (the IMC panel lacks a PD-1 antibody). That this checkpoint trajectory reproduces across independent cohorts, molecular layers, and species argues against a platform-specific artifact and instead supports a genuine biological transition: transient checkpoint upregulation during the Hyperplasia-to-EarlyAdenoma window, superseded by the durable myeloid-dominated immunosuppression described above once lesions become invasive. These tiers are complementary, non-pooled support rather than a matched validation of the same patients; as in the original study on this cohort, no patient-matched human scRNA-seq exists. This reproducible checkpoint signal lends independent support to the jamming-physics interpretation above, in which rising immune infiltration and checkpoint-mediated immune quenching are proposed as the biological driver lowering effective intercellular adhesion and precipitating the fragmentation transition.

These stage-locked vulnerabilities challenge one-size-fits-all checkpoint inhibition strategies and highlight discrete windows, particularly the AAH–MIA interval, for precise immunoprevention through Treg modulation or myeloid remodeling (Fig. 6; Supplementary Figs. S6–S10, S12). We employed a timed Petri net formalism rather than single-cell trajectory inference methods such as Monocle or PAGA, since our data comprise discrete, pathology-defined population states measured at the protein level across cross-sectional specimens rather than a continuous transcriptomic manifold; timed Petri nets quantify directional population flux between defined stages without requiring pseudotime assumptions or single-cell resolution.

Beyond LUAD, ROSIE’s ability to infer protein-level immune landscapes directly from ubiquitous H&E slides positions it as a broadly applicable platform for spatial biomarker discovery across precursor settings, including colorectal adenomas, Barrett’s esophagus, and pancreatic intraepithelial neoplasia. What generalizes is not the LUAD-specific findings themselves but the pipeline that produced them: cohort-scale virtual multiplexing, DBSCAN-based architectural quantification, and timed Petri net population modeling require only routine archival H&E slides and a pathology-defined precursor continuum, both already available for these other premalignant settings, though specific stage boundaries and cellular players would need to be re-derived rather than assumed. The two-phase immune cascade identified here provides a mechanistic rationale for stage-specific therapeutic targeting: early adaptive immune collapse at the AAH–AIS boundary suggests a window for immune checkpoint intervention, whereas the myeloid-dominant landscape of MIA–IAC highlights potential for myeloid reprogramming strategies. Coupling these signatures with prospective longitudinal sampling and multimodal profiling will be essential to establish causal mechanisms and validate clinical utility.

## Conclusion

Architectural fragmentation emerges as a structural inflection point, marking the shift from cohesive, immune-surveilled precursors to disordered, immunosuppressed lesions. The conserved two-phase immune cascade, early adaptive engagement followed by myeloid-driven suppression, reveals discrete stage-locked vulnerabilities in the evolving microenvironment. By deriving these spatial features directly from routine H&E slides, this framework provides a scalable path for integrating spatial biomarkers into clinical workflows, thereby enabling earlier detection, improved risk stratification, and precision interception. Coupling these structural and immune signatures with prospective sampling and multimodal profiling will further refine our understanding of the early malignant transition and accelerate the development of preventive strategies.

## Acknowledgements

We thank the Texas Advanced Computing Center (TACC) for providing computational resources through the project “Predictions of Multitargeting Drug Response for Colorectal Cancer Using AI Techniques and Single-Cell Analysis” (Grant No. MCB23032). We acknowledge the support provided by the Texas Workforce Commission.

## Funding

This work was supported by the National Cancer Institute of the National Institutes of Health under Award Number R01CA234629-01 and the AACR-Johnson & Johnson Lung Cancer Innovation Science Grant (18-90-52-ZHAN).

## Data Availability

H&E-stained whole-slide images and associated clinical metadata cannot be deposited in public repositories owing to patient privacy regulations and institutional data governance restrictions. Access to original high-resolution WSIs may be granted to qualified researchers for non-commercial research purposes upon request from the corresponding author, requiring a formal Data Use Agreement with The University of Texas MD Anderson Cancer Center. All de-identified, analysis-ready feature matrices, including single-cell biomarker predictions, phenotypic labels, Cartesian spatial coordinates, and niche-level summaries, are publicly available at https://github.com/bayjuan5/LUAD-spatial-atlas-A versioned archive will also be deposited on Zenodo upon acceptance.

## Code Availability

All computational analyses were performed in Python (v3.9+) and R (v4.1+) in a version-controlled Linux environment. All analysis codes are openly available at https://github.com/bayjuan5/LUAD-spatial-atlas-, and the repository includes complete scripts, environment specifications, and documentation to facilitate implementation of the ROSIE-enabled spatial profiling framework in independent cohorts.

## Ethics Approval and Consent to Participate

This study was conducted in accordance with the Declaration of Helsinki and approved by the Institutional Review Board (IRB) of The University of Texas MD Anderson Cancer Center. All human tissue samples were obtained from archival pathology repositories by using IRB-approved protocols. Informed consent was waived by the IRB owing to the retrospective nature of the study and the use of de-identified archival specimens.

## Author Contribution Statement

BH conceived the study, designed the computational framework, performed all analyses, and drafted the manuscript. BZ performed wetlab experimental validation, contributed clinical sample curation, and participated in manuscript writing and revision. Both authors reviewed and approved the final manuscript.

## Declaration of Competing Interest

The authors declare no competing interests.

## Supplementary Information

**Figure S1.**
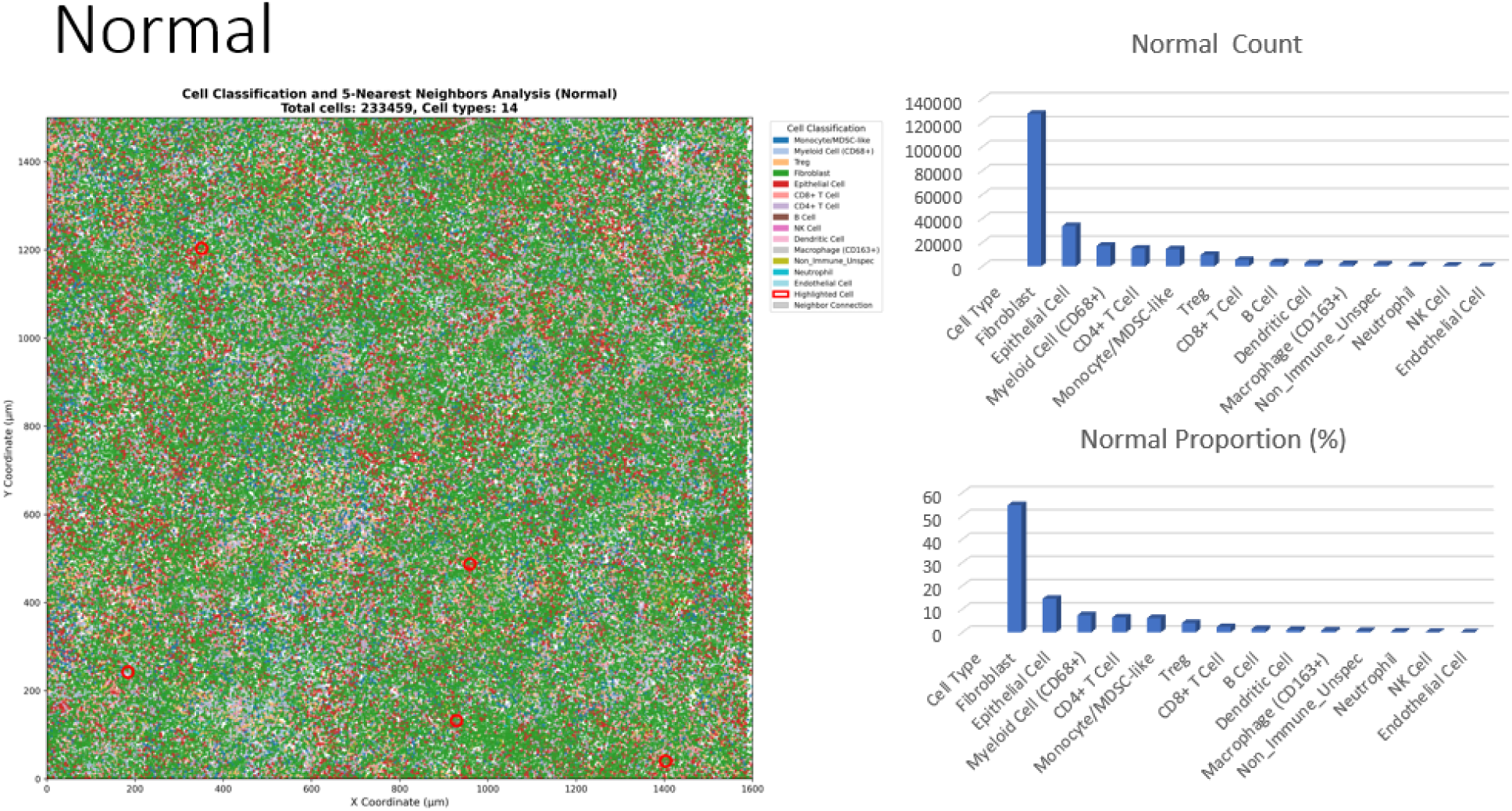
Spatial distribution and quantification of cell types in normal lung tissue. Spatial cell classification map of 233,459 cells across 14 cell types in normal lung parenchyma. Left: Each point represents a single cell positioned by its X–Y coordinates and colored by predicted cell identity. Normal tissue exhibits a cohesive architecture dominated by epithelial and endothelial compartments, with immune populations dispersed throughout the region. Right: Bar plots summarizing absolute cell counts (top) and relative proportions (bottom). Epithelial cells constitute the largest population (∼140,000 cells; ∼60%), followed by endothelial cells and fibroblasts. Immune subsets—including CD8⁺ T cells, CD4⁺ T cells, Tregs, B cells, NK cells, dendritic cells, monocyte/MDSC-like cells, macrophages (CD163⁺), neutrophils, and myeloid CD68⁺ cells—appear at lower but detectable frequencies. The 5-nearest-neighbors graph embedded in the spatial map highlights local micro-architectural organization and baseline cell–cell proximity patterns characteristic of normal lung tissue. These features establish the reference framework for comparing microenvironmental remodeling across pre-malignant and malignant stages of LUAD progression.

**Fig S2.**
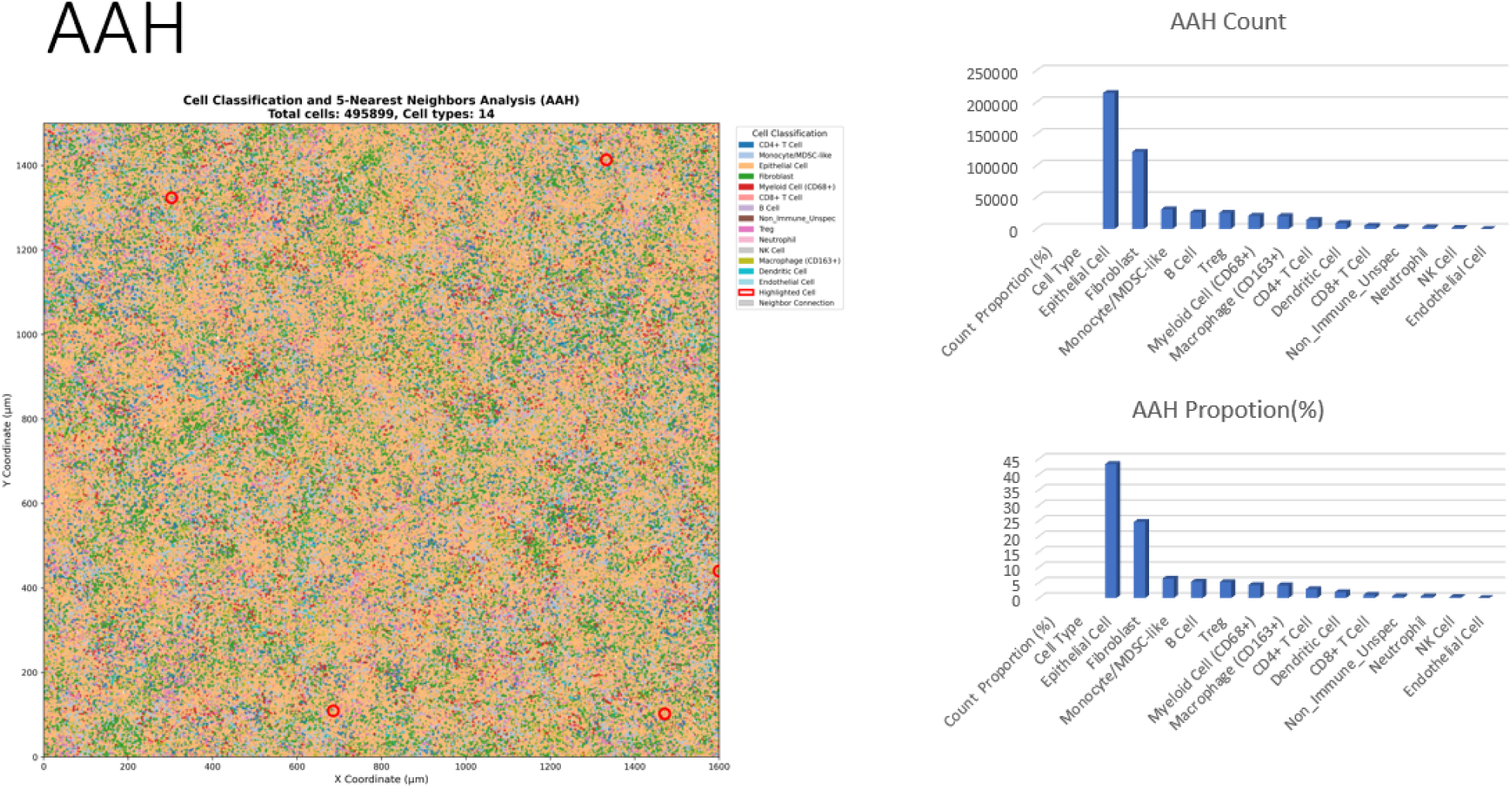
Spatial distribution and quantification of cell types in Atypical Adenomatous Hyperplasia (AAH) Spatial cell classification map of 511,157 cells across 14 cell types in AAH lesions, representing early pre-malignant stages of lung adenocarcinoma. Left: Each point represents a single cell positioned by its X–Y coordinates and colored by predicted cell identity. Red circles highlight regions used for 5-nearest-neighbors analysis. Compared with normal lung tissue, AAH exhibits increased spatial heterogeneity, with emerging clusters of fibroblasts, immune infiltrates, and altered epithelial organization. Right: Bar plots summarizing absolute cell counts (top) and relative proportions (bottom). Epithelial cells remain the predominant population (∼220,000 cells; ∼45%), followed by fibroblasts and endothelial cells. Immune populations—including CD4⁺ and CD8⁺ T cells, B cells, monocyte/MDSC-like cells, macrophages (CD163⁺), dendritic cells, neutrophils, NK cells, and Tregs—show increased representation relative to normal tissue. This shift toward greater cellular diversity and immune infiltration reflects the transitional microenvironment characteristic of AAH, marking the earliest detectable remodeling events along the LUAD precursor continuum.

**Fig S3.**
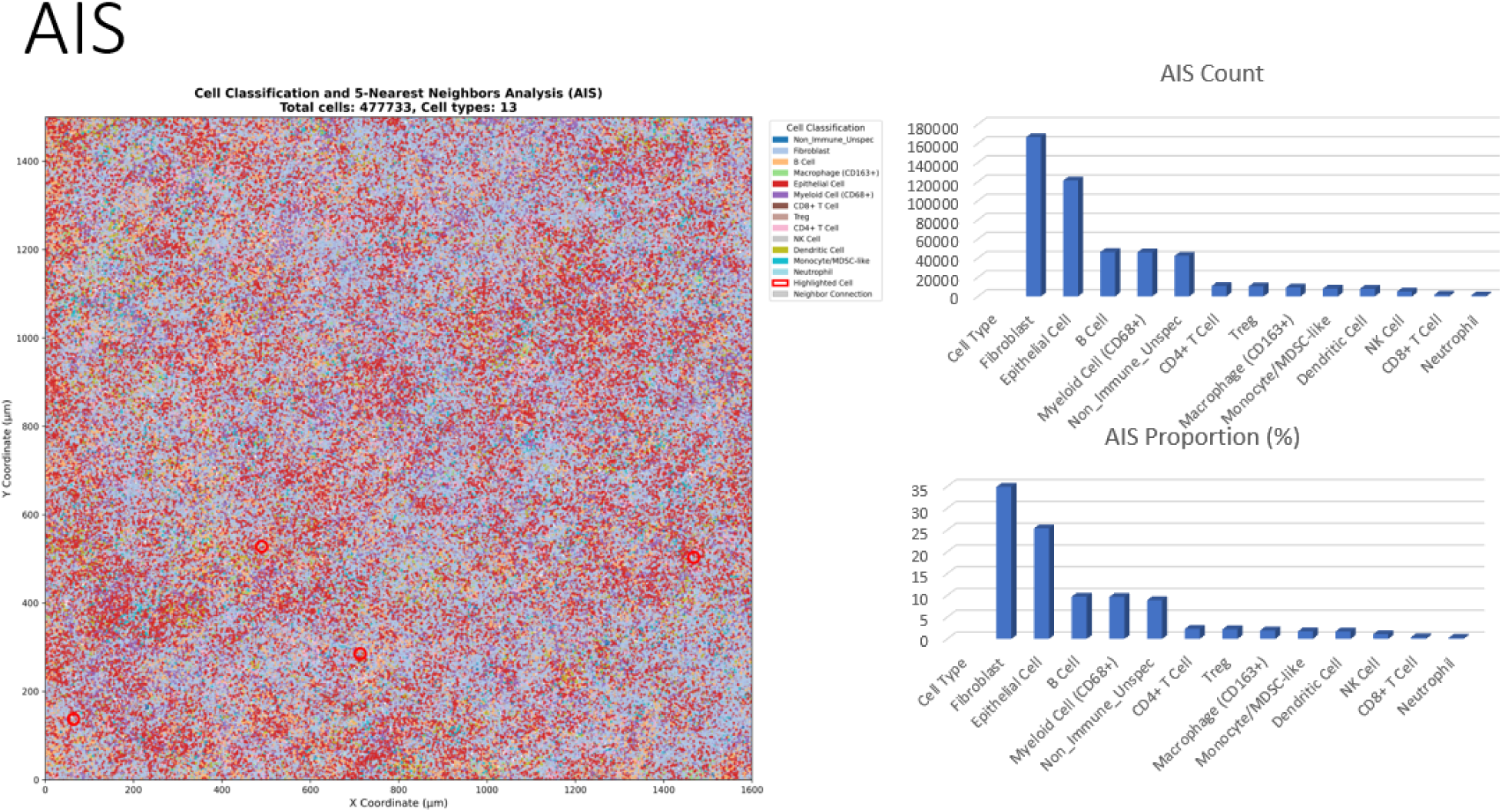
Spatial distribution and quantification of cell types in Adenocarcinoma in Situ (AIS) Spatial cell classification map of **437,918 cells** across **14 cell types** in AIS lesions, representing a more advanced pre-malignant stage in lung adenocarcinoma progression. Left: Each point represents a single cell positioned by its X–Y coordinates and colored by predicted cell identity. Red circles highlight regions used for **5-nearest-neighbors** analysis. The AIS architecture shows **pronounced spatial clustering**, with red-dominant regions interspersed within a blue–purple background, reflecting increased segregation of epithelial, fibroblast, and immune compartments compared with earlier stages. Right: Bar plots summarizing **absolute cell count** (top) and **relative proportions** (bottom). **Epithelial cells** remain the predominant population (∼175,000 cells; ∼35%), followed by **fibroblasts** (∼27%) and **endothelial cells**. Immune populations—including B cells, CD4⁺ and CD8⁺ T cells, Tregs, monocyte/MDSC-like cells, macrophages (CD163⁺), dendritic cells, NK cells, and neutrophils—show **further expansion** relative to AAH. The marked spatial segregation and shifting stromal–immune composition reflect the **progressive remodeling of the tumor microenvironment** as lesions advance from AAH to AIS along the LUAD precursor continuum.

**Fig S4.**
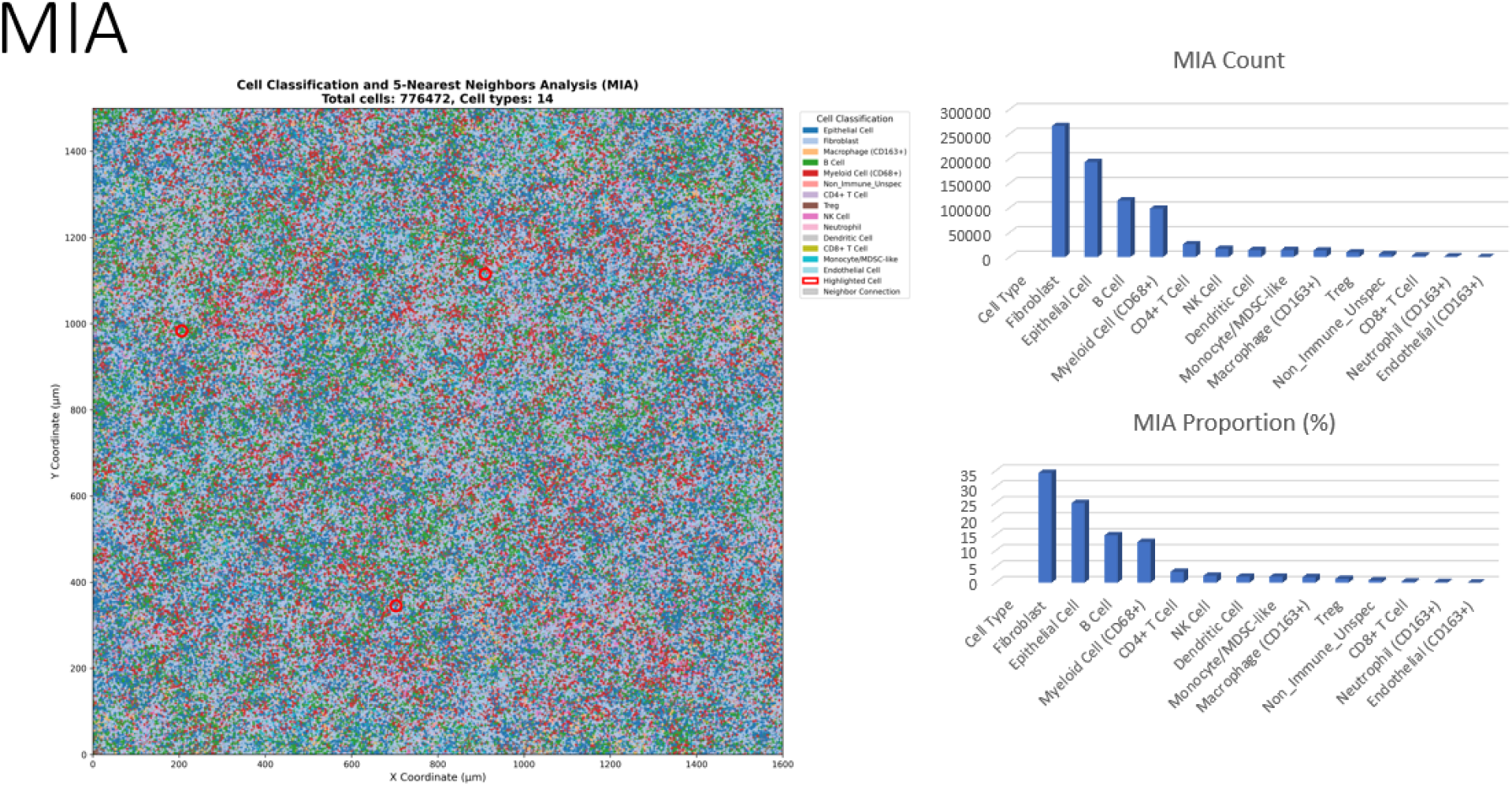
Spatial distribution and quantification of cell types in Minimally Invasive Adenocarcinoma (MIA) Spatial cell classification map of 59,027 cells (representative subsample) across 14 cell types in MIA tissue. Left: Each point represents a single cell positioned by its X–Y coordinates and colored by predicted cell identity. The spatial architecture shows increased cellular diversification compared with AIS, with emerging stromal and immune niches characteristic of early invasion. Right: Bar plots summarizing absolute cell counts for each cell type. B cells constitute the most abundant population (21,982 cells), followed by fibroblasts (16,077 cells) and myeloid CD68⁺ cells (10,334 cells). Regulatory T cells (6,215 cells) and monocyte/MDSC-like cells (5,766 cells) show notable expansion relative to AIS. Additional populations—including CD4⁺ T cells, epithelial cells, neutrophils, macrophages (CD163⁺), dendritic cells, NK cells, CD8⁺ T cells, and non-immune unspecified cells—are present at lower frequencies. This cellular composition reflects the MIA stage, where early stromal invasion begins, marked by substantial B-cell infiltration, reorganization of stromal components, and a more complex immune microenvironment compared with the CD4⁺ T-cell–dominated AIS stage.

**Fig S5.**
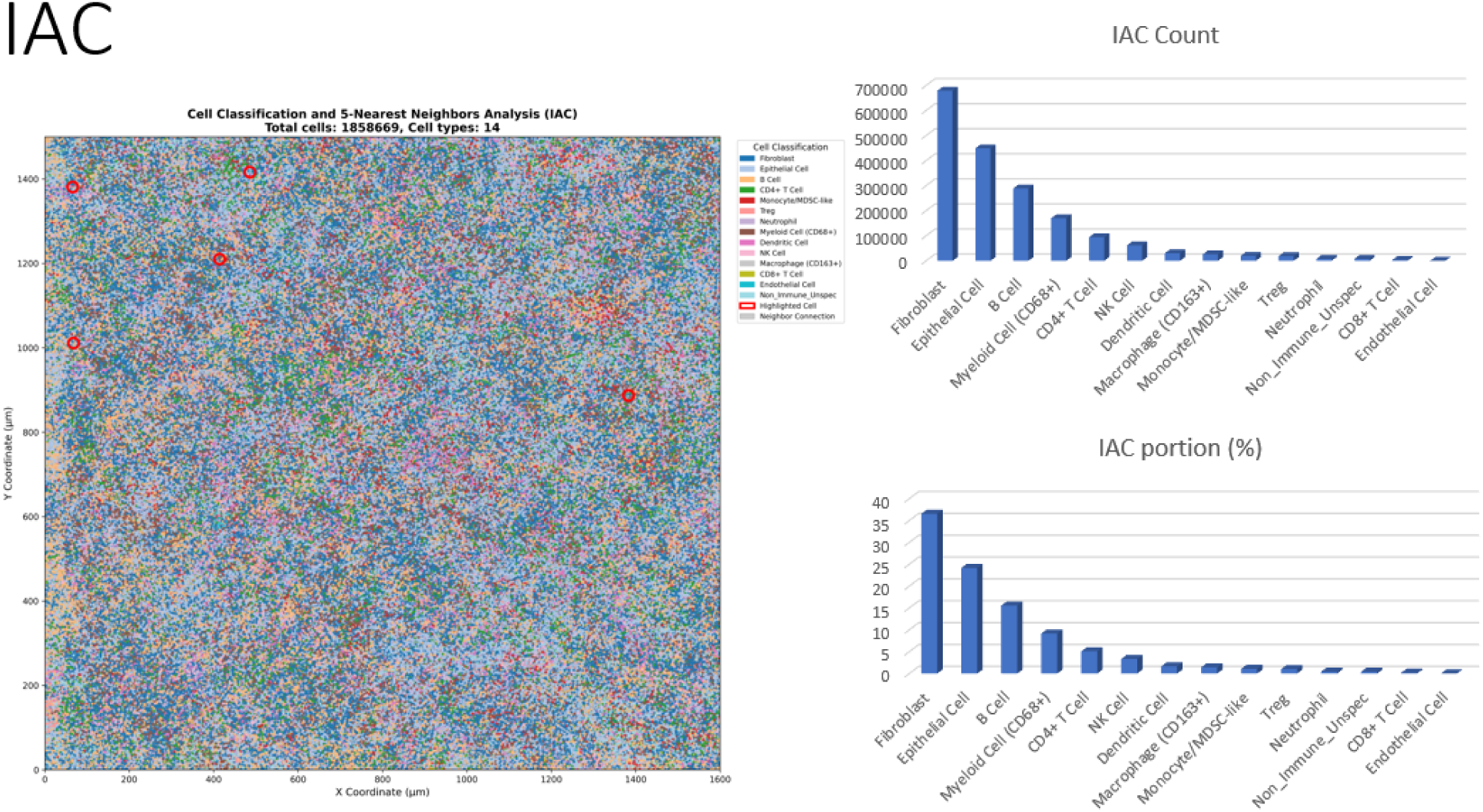
Spatial distribution and quantification of cell types in Invasive Adenocarcinoma (IAC) Spatial cell classification map of **1,858,669 cells** across **14 cell types** in IAC tissue, representing fully invasive lung adenocarcinoma. Left: Each point represents a single cell positioned by its X–Y coordinates and colored by predicted cell identity. Red circles highlight regions used for **5-nearest-neighbors** analysis. The spatial architecture displays **extensive cellular heterogeneity**, with complex intermixing of epithelial, stromal, and immune populations forming a highly diverse mosaic characteristic of invasive cancer. Right: Bar plots summarizing **absolute cell counts** (top) and **relative proportions** (bottom). **Fibroblasts** constitute the dominant population (∼700,000 cells; ∼38%), reflecting a dramatic expansion of the stromal compartment compared with earlier stages. **Epithelial cells** (∼26%) and **B cells** (∼17%) represent the next most abundant populations. The invasive stage also exhibits **robust immune infiltration**, with substantial representation of myeloid CD68⁺ cells, CD4⁺ T cells, NK cells, dendritic cells, macrophages (CD163⁺), monocyte/MDSC-like cells, neutrophils, Tregs, CD8⁺ T cells, non-immune/unspecified cells, and endothelial cells. This marked stromal expansion, combined with diverse immune infiltration and loss of organized tissue architecture, reflects the **complex tumor microenvironment of fully invasive adenocarcinoma**, representing the culmination of progressive cellular and structural remodeling across the LUAD continuum.

**Fig S6.**
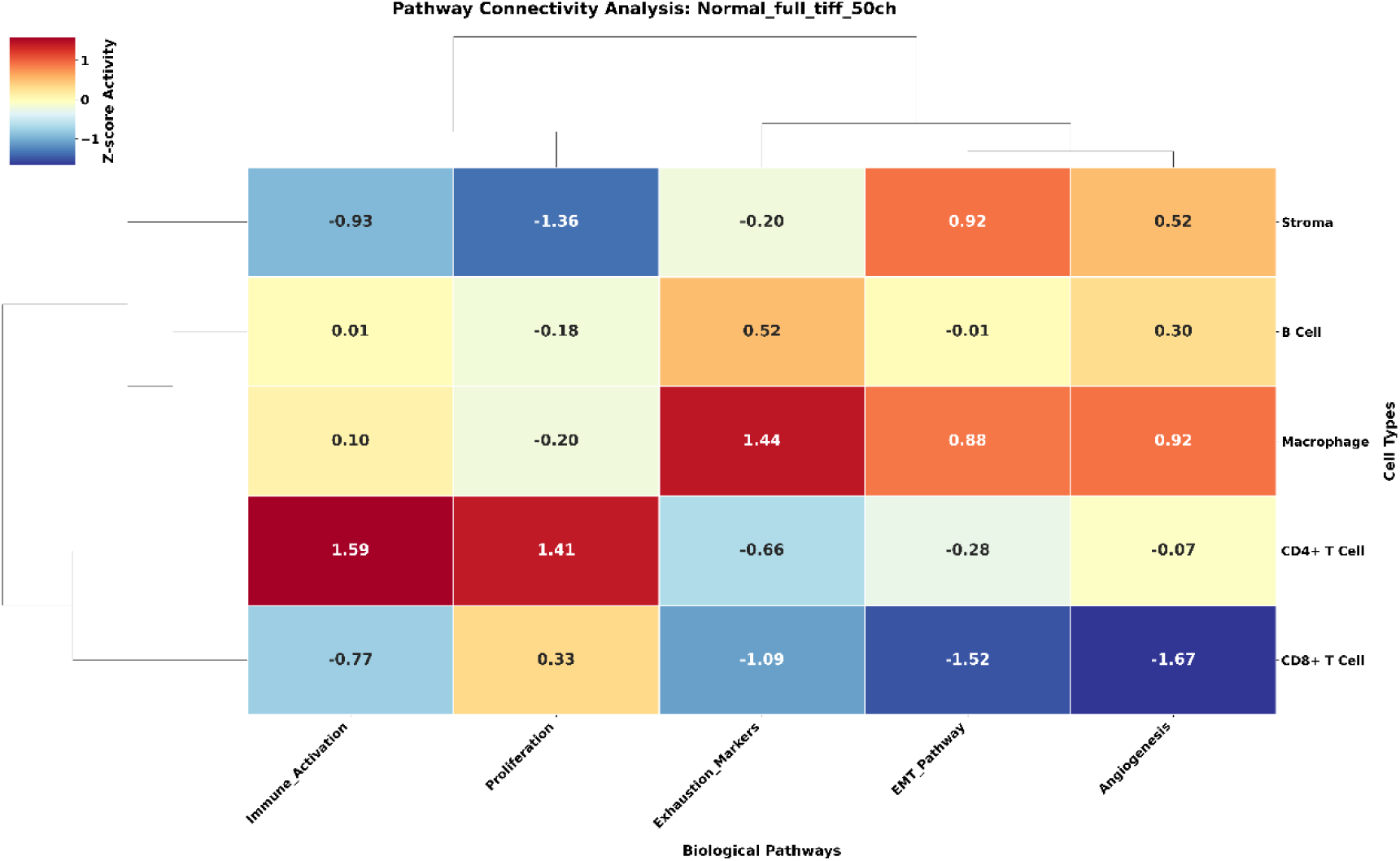
Niche-specific pathway activity in Normal lung tissue. Heatmap showing Z-score–normalized pathway activity across major cell types in Normal lung tissue. Rows represent cell types and columns represent biological pathways, including Immune Activation, Proliferation, Exhaustion Markers, EMT, and Angiogenesis. Normal tissue displays uniformly elevated stromal and epithelial pathway activity with balanced immune signaling, reflecting a homeostatic microenvironment prior to neoplastic transformation.

**Fig S7.**
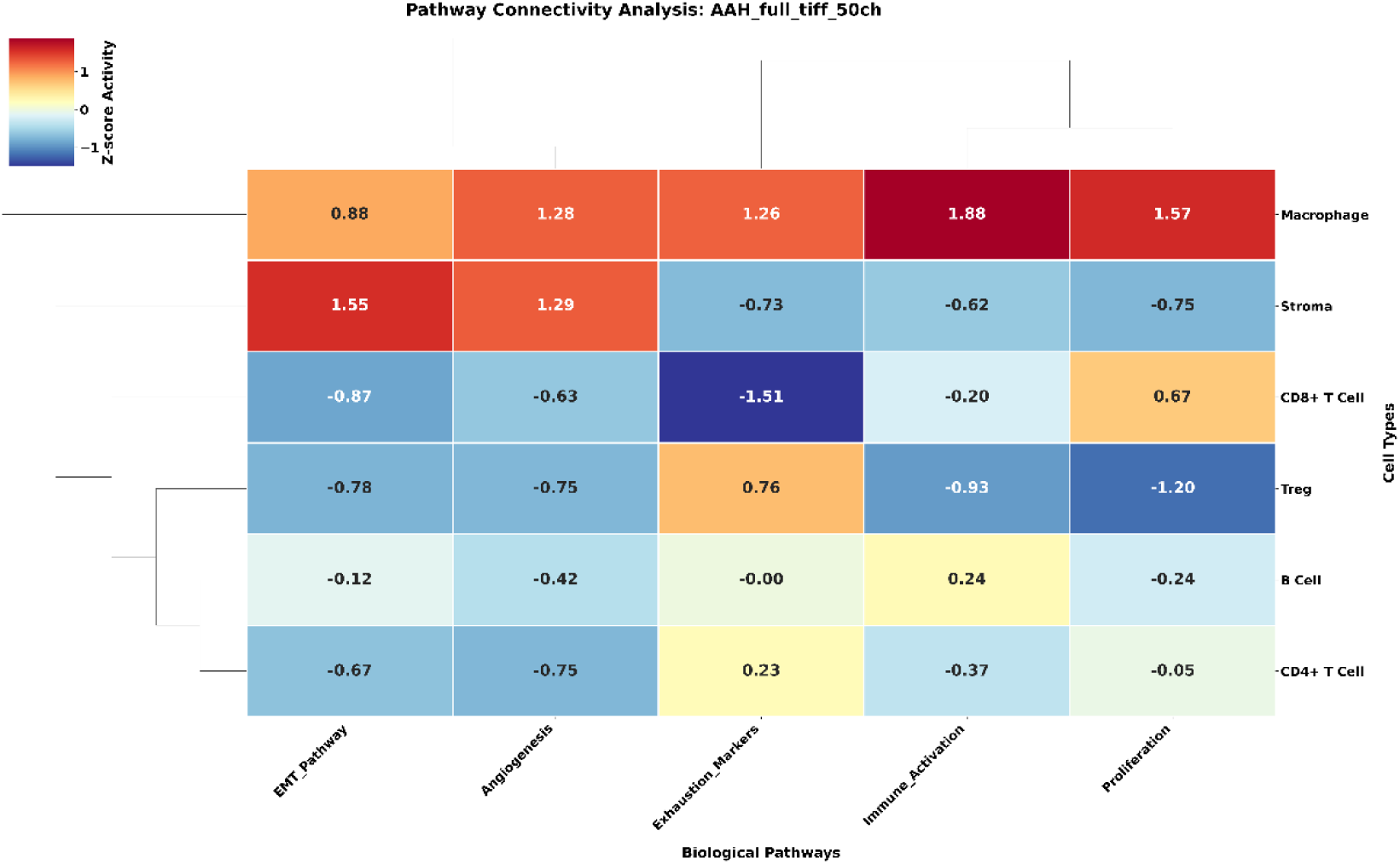
Niche-specific pathway activity in Atypical Adenomatous Hyperplasia (AAH). Heatmap depicting pathway activity across cellular niches in AAH lesions. Early deviations from homeostasis emerge, including increased proliferative and immune-activation signals in epithelial and macrophage compartments. These early pathway shifts mark the onset of microenvironmental remodeling associated with pre-neoplastic change.

**Fig S8.**
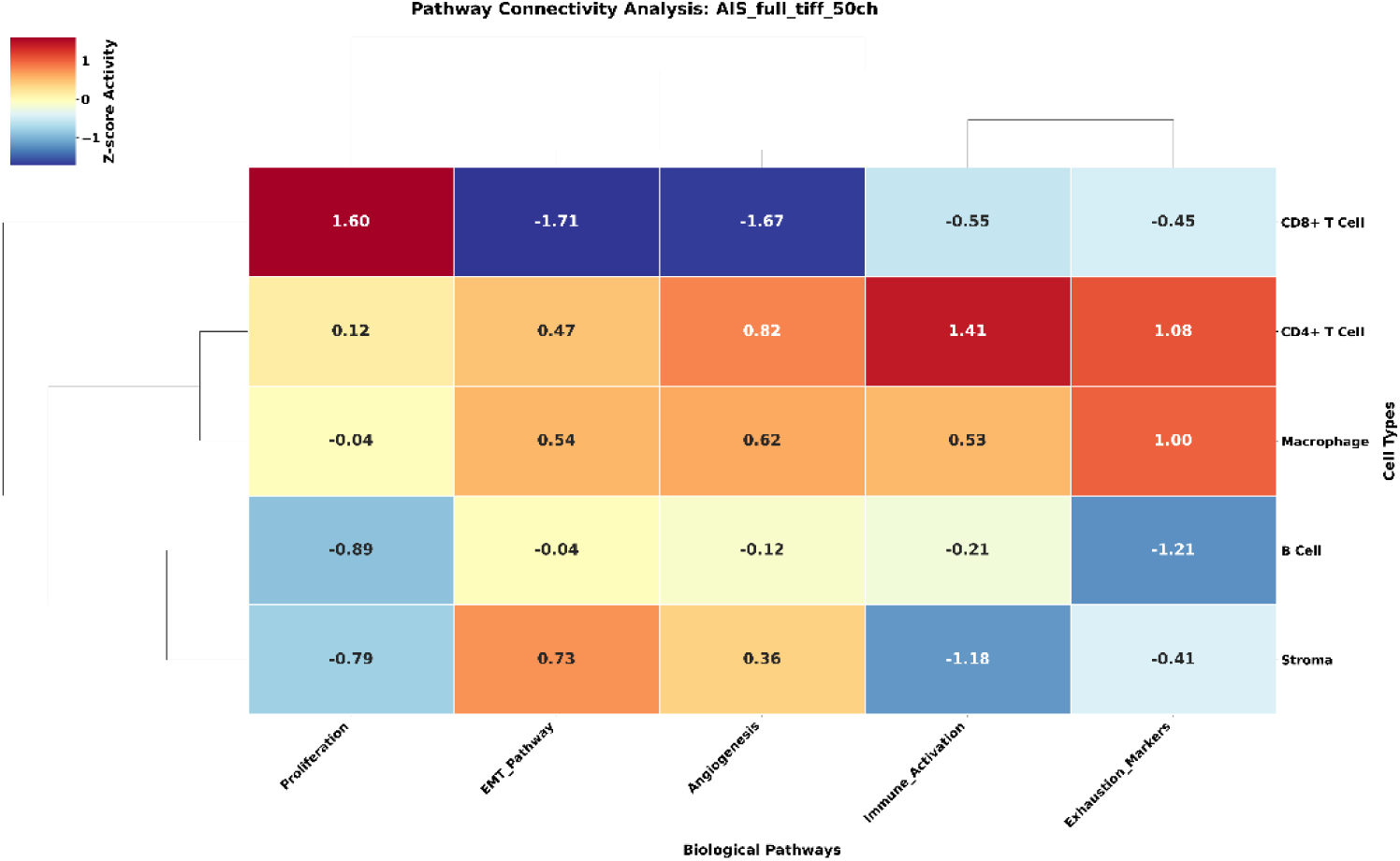
Niche-specific pathway activity in Adenocarcinoma in Situ (AIS). Heatmap illustrating pathway activity across cell types in AIS. A coordinated decline in immune activation and stromal signaling becomes evident across multiple niches, while epithelial proliferative programs intensify. These changes reflect the establishment of a less immunogenic, more epithelial-dominant microenvironment characteristic of in situ malignancy.

**Fig S9.**
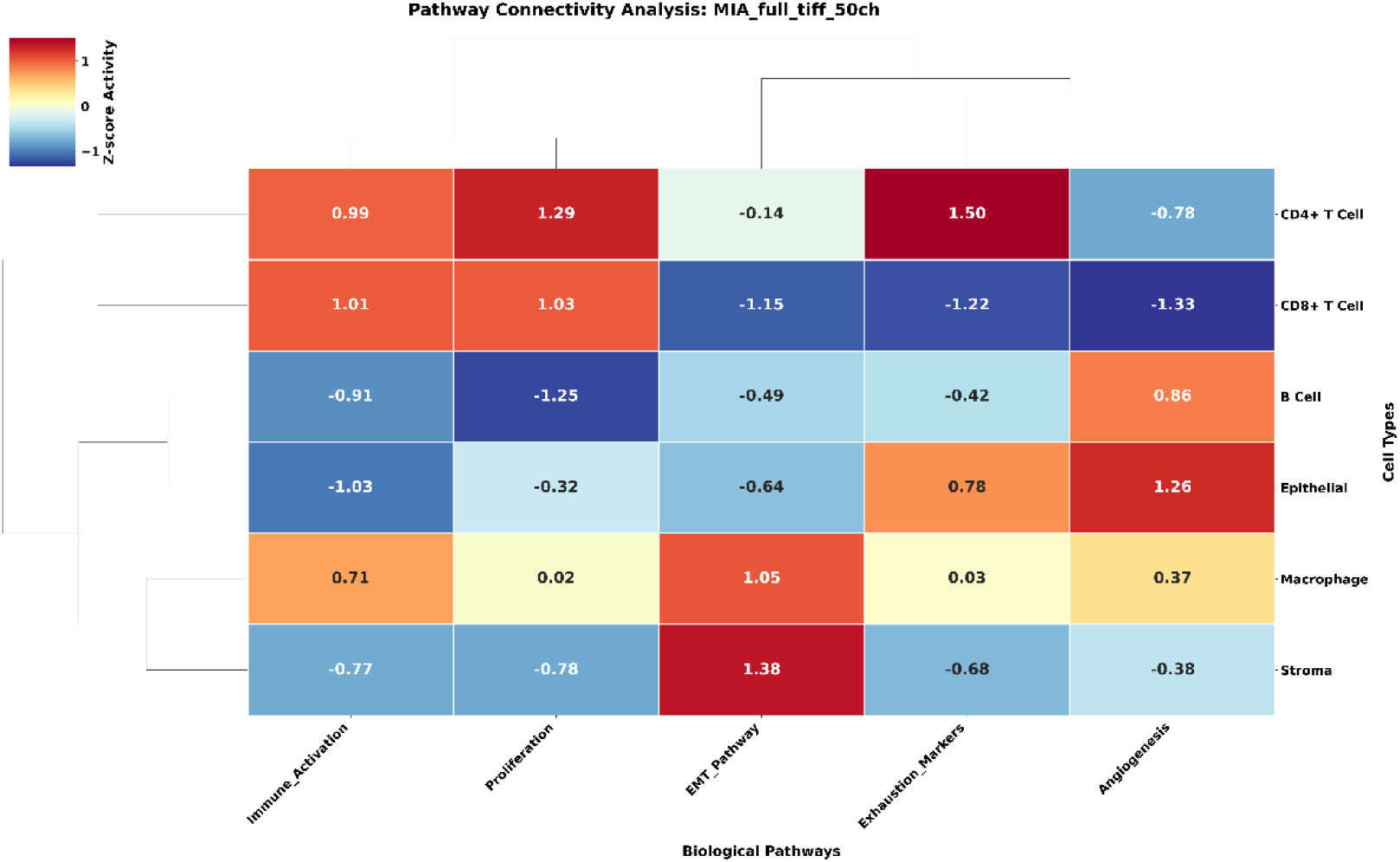
Niche-specific pathway activity in Minimally Invasive Adenocarcinoma (MIA). Heatmap showing pathway activity across cellular niches in MIA. Immune and stromal suppression deepens, particularly within myeloid and fibroblast compartments, while epithelial cells exhibit heightened proliferative and EMT-associated activity. These patterns indicate progressive immune evasion and early invasive potential.

**Fig S10.**
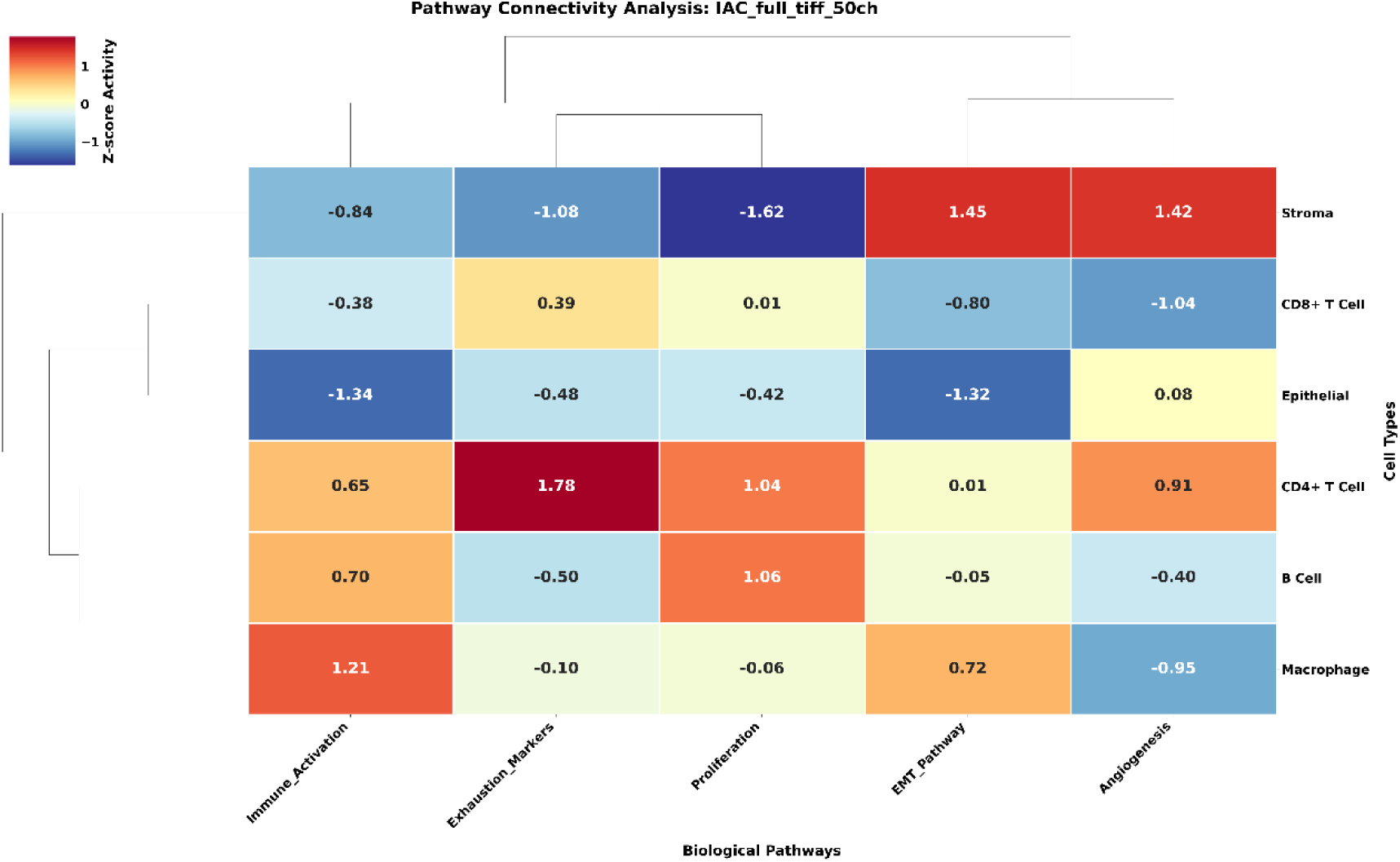
Niche-specific pathway activity in Invasive Adenocarcinoma (IAC). Heatmap summarizing pathway activity across cell types in IAC. Nearly all immune and stromal niches exhibit marked suppression of immune activation and checkpoint pathways, whereas malignant epithelial cells display dominant proliferative and angiogenic activity. This niche-level reprogramming reflects a fully established tumor-permissive microenvironment supporting invasive growth.

**Fig S11.**
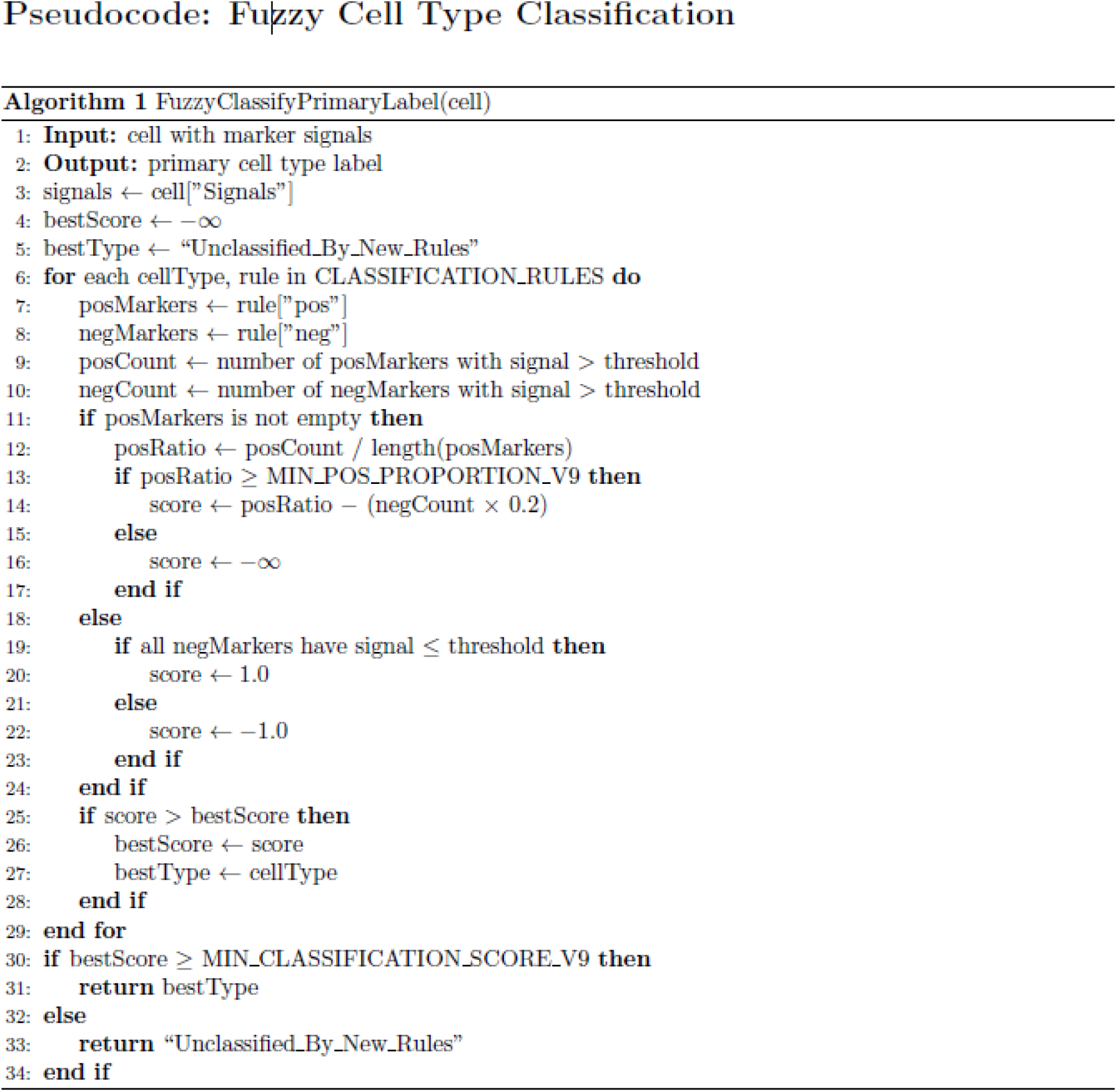
Pseudocode for fuzzy cell type classification algorithm. This figure outlines the logic of the FuzzyClassifyPrimaryLabel algorithm used to assign primary cell type labels based on multiplexed marker signals. The algorithm iteratively evaluates each candidate cell type defined in the classification rule set, computing a fuzzy score based on the proportion of positive markers detected and penalizing the presence of negative markers. Cells are assigned the label with the highest score exceeding a minimum classification threshold; otherwise, they are marked as Unclassified_By_New_Rules. This approach enables robust and interpretable annotation of diverse immune, stromal, and epithelial populations in LUAD tissue samples.

**Table S2.** Marker-based cell classification rules used for fuzzy labeling. This table defines the positive and negative marker sets used to assign primary cell type labels via the fuzzy classification algorithm. Each cell type is characterized by canonical marker combinations, with positive markers required for identity confirmation and negative markers used to exclude conflicting phenotypes. The rule set spans immune subsets (e.g., Tregs, CD4⁺/CD8⁺ T cells, B cells, NK cells), myeloid populations (e.g., macrophages, dendritic cells, monocytes), and structural cells (e.g., epithelial, fibroblast, endothelial), ensuring biologically interpretable and reproducible annotation across LUAD tissue samples.

| Cell Type | Positive Markers | Negative Markers |
| --- | --- | --- |
| Treg | CD45, CD3e, CD4, FoxP3 | CD8, CD20, CD68, PanCK |
| CD8+ T Cell | CD45, CD3e, CD8 | CD4, FoxP3, CD20, CD68, PanCK |
| CD4+ T Cell | CD45, CD3e, CD4 | CD8, FoxP3, CD20, CD68, PanCK |
| B Cell | CD45, CD20 | CD3e, CD68, CD163, CD66, CD11c, PanCK |
| NK Cell | CD45 | CD3e, CD20, CD68, CD163, CD66, CD11c, PanCK, CD31 |
| Neutrophil | CD45, CD66 | CD3e, CD20, CD68, CD163, CD11c, PanCK |
| Macrophage (CD163+) | CD45, CD163 | CD3e, CD20, CD66, CD11c, PanCK |
| Myeloid Cell (CD68+) | CD45, CD68 | CD3e, CD20, CD66, CD163, CD11c, PanCK |
| Dendritic Cell | CD45, CD11c | CD3e, CD20, CD66, CD68, CD163, PanCK |
| Monocyte/MDSC-like | CD45, CD68 | CD3e, CD20, CD31, PanCK, Vimentin |
| Epithelial Cell | PanCK | CD45, Vimentin |
| Fibroblast | Vimentin | CD45, PanCK |
| Endothelial Cell | CD31 | CD45, PanCK, Vimentin |
| Non-Immune Unspec | — | CD45, PanCK, Vimentin, CD31 |
| Background/Junk | — | CD45, PanCK, Vimentin, CD31 |

**Supplementary Fig. S12.**
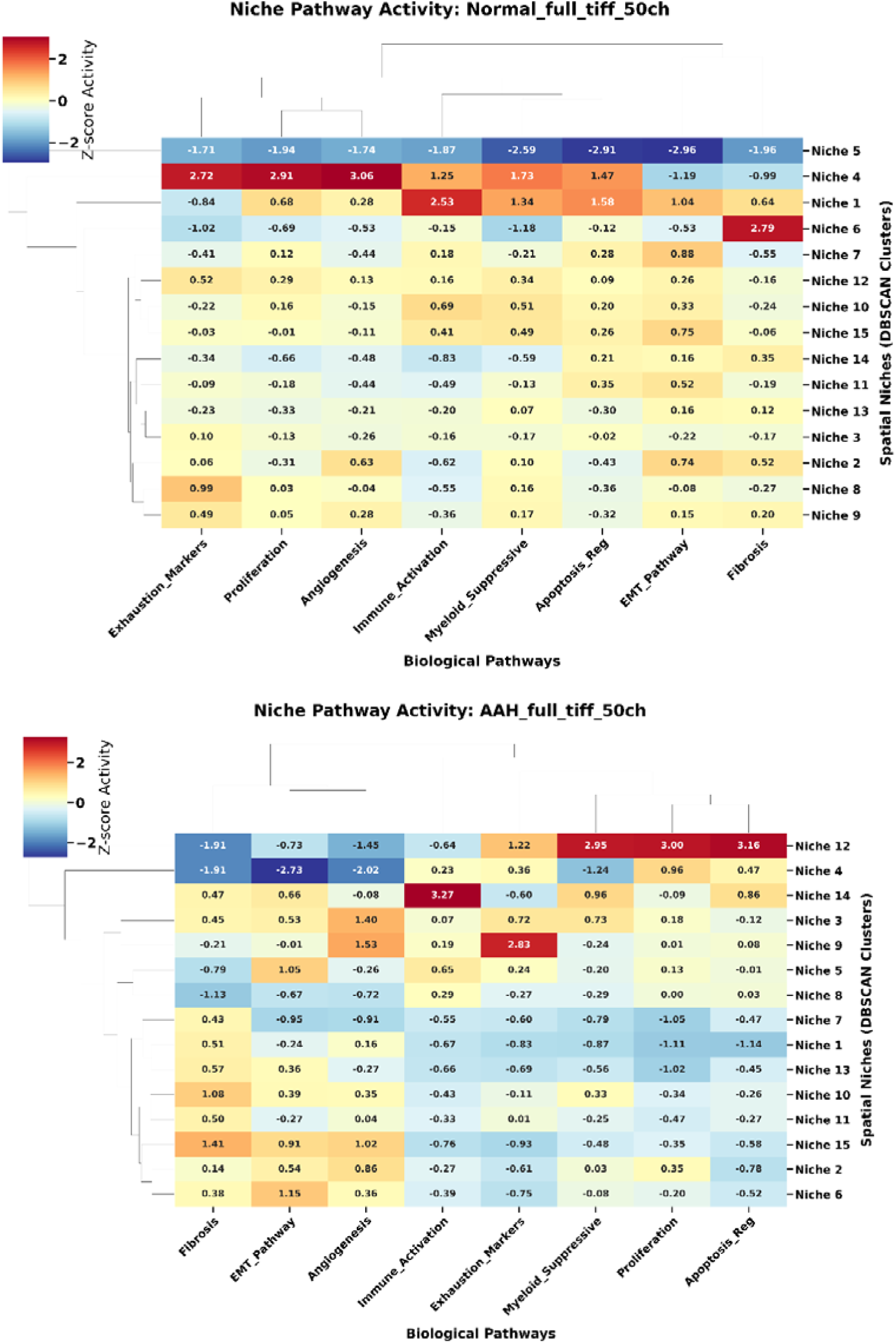
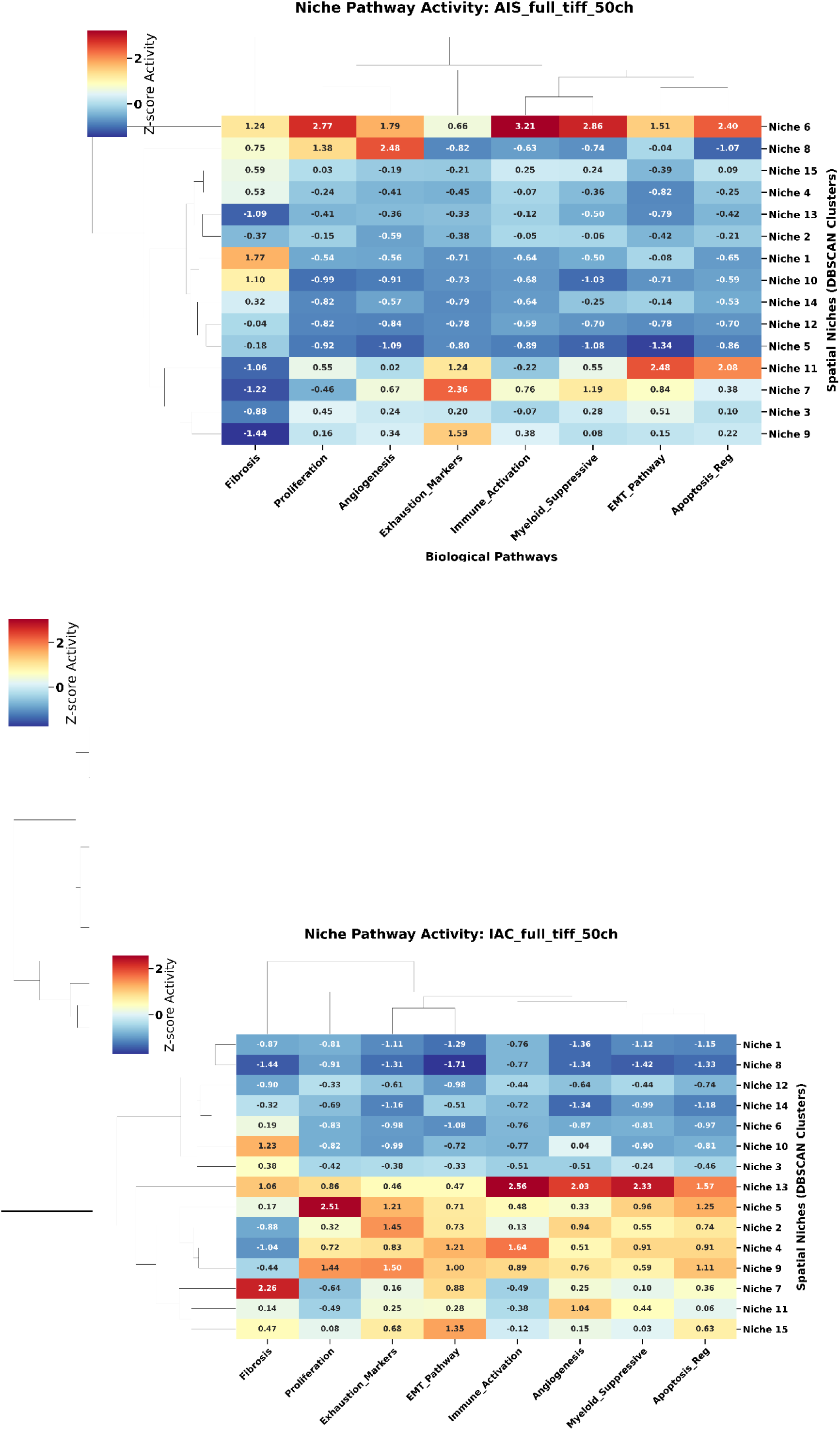
Niche-specific pathway activity across LUAD progression. (a–e) Heatmaps of niche-specific pathway activity at Normal, AAH, AIS, MIA, and IAC stages, respectively. Progressive immunosuppression and stromal quenching are coupled with intensifying epithelial proliferation across stages.

**Supplementary Fig. S13.**
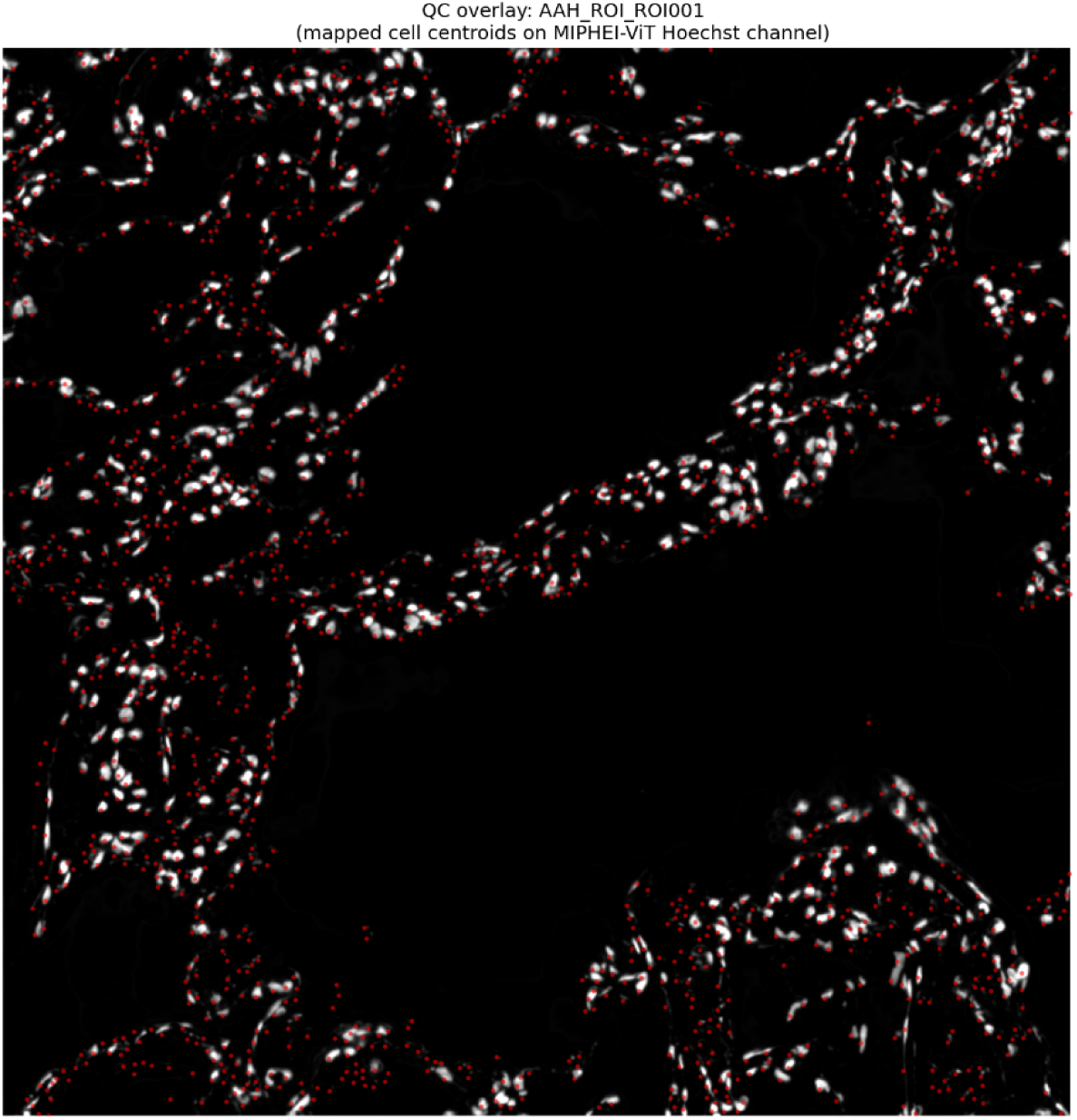
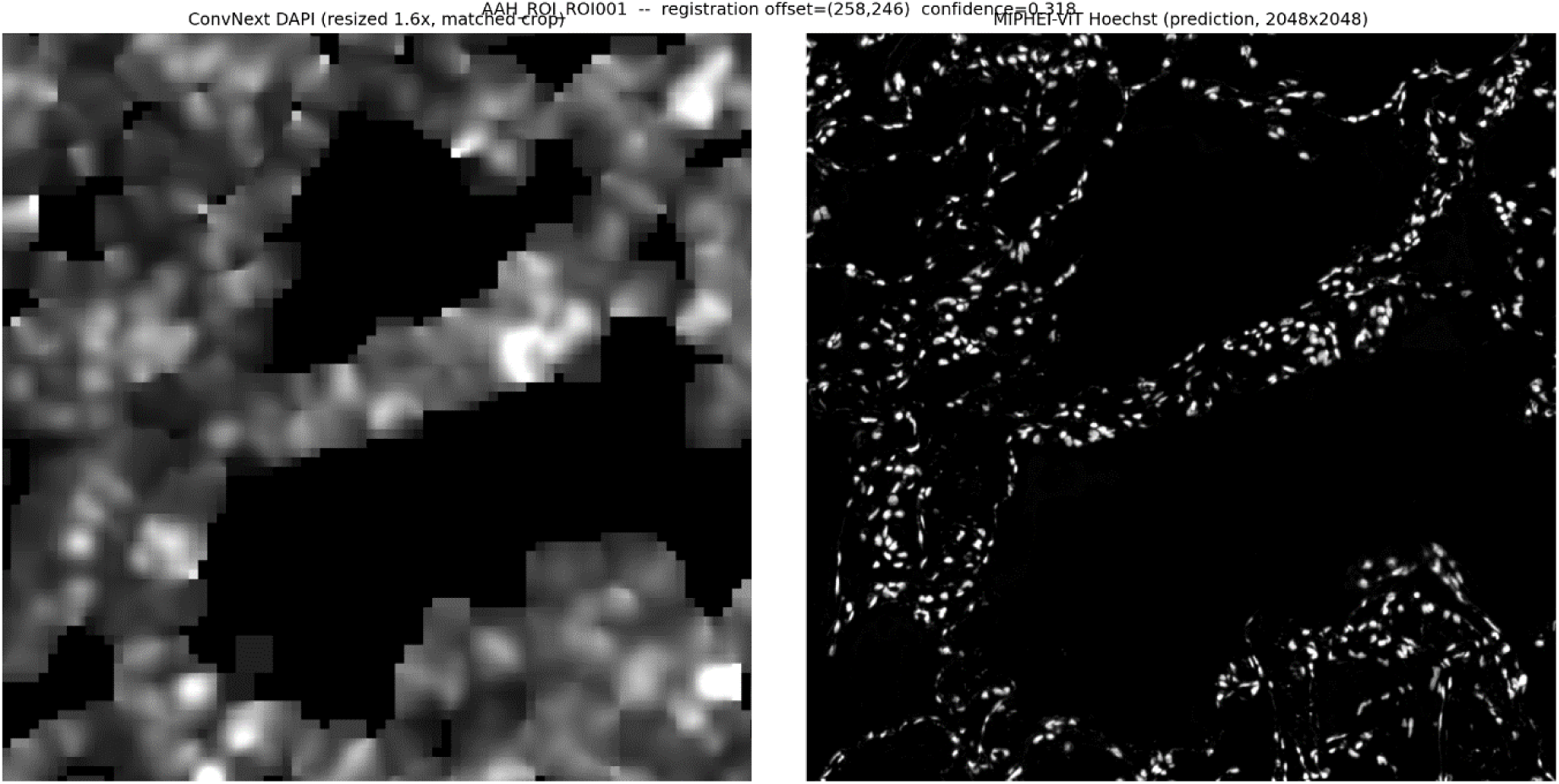
Coordinate registration and resolution comparison between ROSIE and MIPHEI-ViT. (a) Registration QC: mapped cell centroids confirm accurate spatial alignment after per-ROI DAPI–Hoechst template matching (mean registration confidence 0.40, range 0.15–0.56 across 52 ROIs). (b) Side-by-side comparison of ROSIE’s DAPI channel and MIPHEI-ViT’s Hoechst channel for the same registered ROI, illustrating that the resolution mismatch between ROSIE’s patch-wise inference (∼64-pixel blocks) and MIPHEI-ViT’s dense per-pixel output, rather than registration error, underlies the improvement in cross-model concordance observed when markers are compared at native patch resolution (Table 2). (a) QC overlay of ROSIE-derived cell centroids (red) mapped onto MIPHEI-ViT’s Hoechst channel for a representative ROI (AAH_ROI_ROI001), following per-ROI DAPI–Hoechst template-matching registration (see Materials and Methods). Accurate spatial co-localization of centroids with nuclear signal confirms registration validity prior to cross-model marker comparison. (b) Matched crop of ROSIE’s DAPI channel (resized 1.6× to MIPHEI-ViT’s pixel grid; left) alongside MIPHEI-ViT’s Hoechst channel (right) for the same ROI. Large-scale tissue architecture aligns closely between the two models, confirming that registration is accurate; the visible texture difference (soft, blocky ∼64-pixel patches from ROSIE vs. sharp individual nuclei from MIPHEI-ViT) reflects the two models’ differing native resolutions, not misalignment, and is the basis for the native-patch-resolution comparison reported in the Results and Table 2.

**Supplementary Fig. S14.**
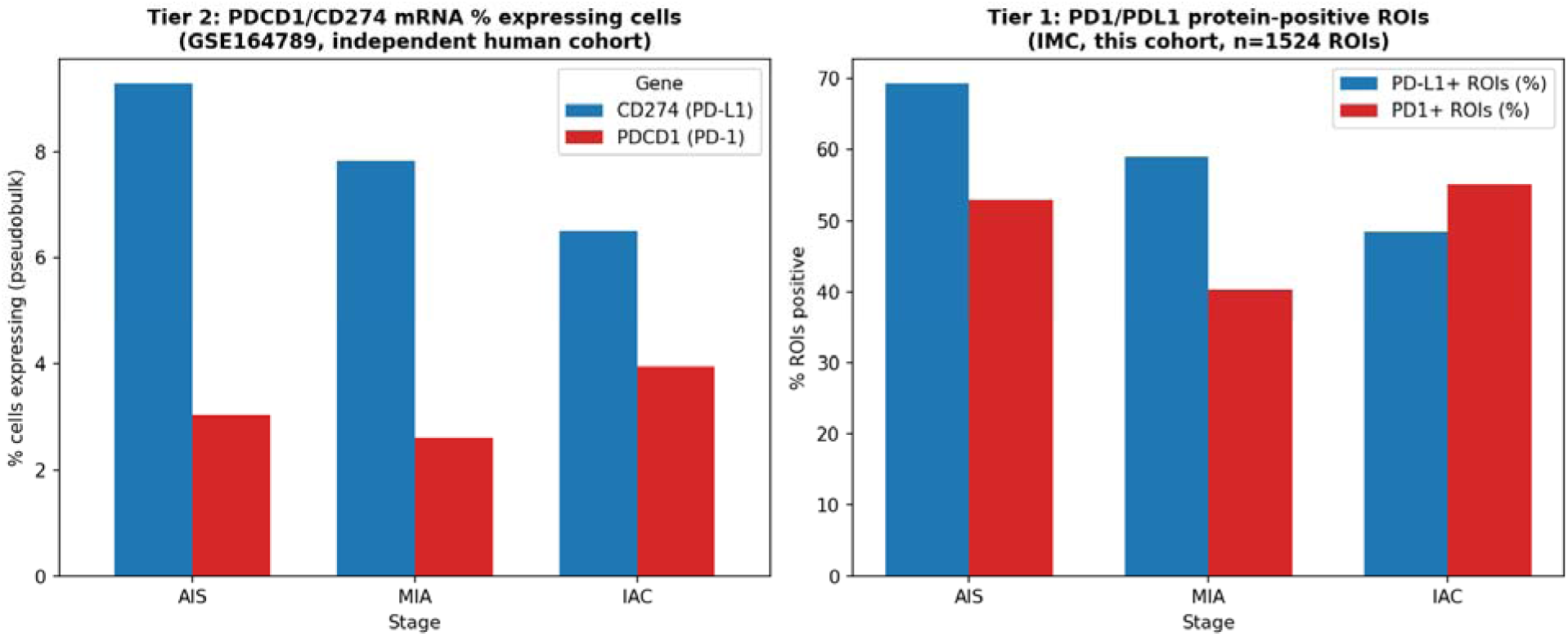
Cross-validation of PD-1/PD-L1 stage trends at the protein and mRNA level. Left: PDCD1/CD274 percent of cells expressing (pseudobulk) across AIS, MIA, and IAC in an independent human scRNA-seq cohort (GSE164789, Tier 2). Right: percent of IMC ROIs classified as PD1-positive or PD-L1-positive across the same three stages in the present 114-specimen cohort (Tier 1, n=1,524 ROIs). PD-L1 declines monotonically at both molecular layers; PD-1 shows a concordant non-monotonic dip at MIA followed by a rebound at IAC in both layers.

**Supplementary Fig. S15.**
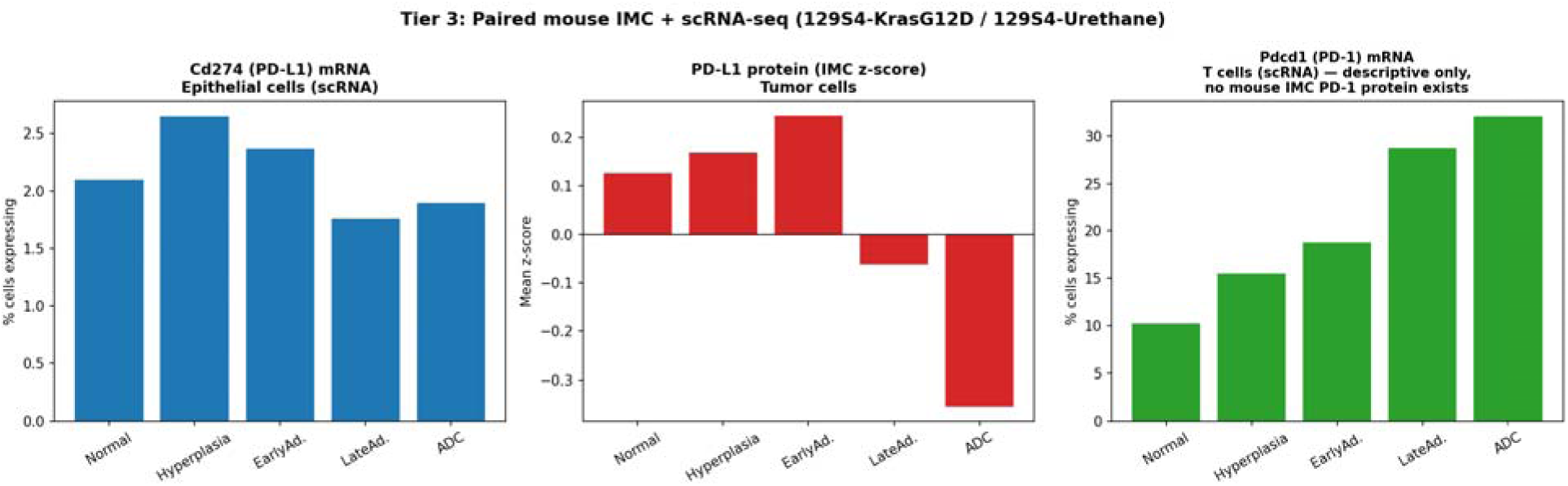
Tier 3 paired protein–mRNA comparison in mouse precancer models. Left: Cd274 (PD-L1) mRNA percent of epithelial cells expressing (scRNA-seq) across the five-stage progression in 129S4-KrasG12D/129S4-Urethane models. Middle: PD-L1 protein (IMC, mean per-ROI z-score) in tumor cells across the same stages and models. Right: Pdcd1 (PD-1) mRNA percent of T cells expressing across the same progression, shown descriptively; no corresponding mouse IMC PD-1 protein measurement exists for cross-validation.

## Notes

### Competing Interest Statement

The authors have declared no competing interest.

### Summary of Updates

This revision updates the manuscript title to “Architectural Fragmentation and Immune Reprogramming Define Early Malignant Transition in Lung Adenocarcinoma.” Major additions since the original preprint include: (1) an independent cross-model validation using MIPHEI-ViT, a second virtual multiplexing framework, performed across 52 regions of interest (203,109 cells) to test whether structural and immune findings generalize beyond ROSIE; (2) expanded validation against imaging mass cytometry ground truth across 25 precursor samples; (3) a jamming-physics interpretation of the architectural fragmentation trajectory; (4) cross-species and cross-molecular-layer validation of the PD-1/PD-L1 checkpoint signal using independent human and mouse datasets; and (5) an updated Discussion and Conclusion reflecting these additional analyses. The corresponding author has been updated to Bo Zhu.

